# Substrate specificity profiling of SARS-CoV-2 main protease enables design of activity-based probes for patient-sample imaging

**DOI:** 10.1101/2020.03.07.981928

**Authors:** Wioletta Rut, Katarzyna Groborz, Linlin Zhang, Xinyuanyuan Sun, Mikolaj Zmudzinski, Bartlomiej Pawlik, Wojciech Młynarski, Rolf Hilgenfeld, Marcin Drag

## Abstract

In December 2019, the first cases of infection with a novel coronavirus, SARS-CoV-2, were diagnosed in Wuhan, China. Due to international travel and human-to-human transmission, the virus spread rapidly inside and outside of China. Currently, there is no effective antiviral treatment for coronavirus disease 2019 (COVID-19); therefore, research efforts are focused on the rapid development of vaccines and antiviral drugs. The SARS-CoV-2 main protease constitutes one of the most attractive antiviral drug targets. To address this emerging problem, we have synthesized a combinatorial library of fluorogenic substrates with glutamine in the P1 position. We used it to determine the substrate preferences of the SARS-CoV and SARS-CoV-2 main proteases, using natural and a large panel of unnatural amino acids. On the basis of these findings, we designed and synthesized an inhibitor and two activity-based probes, for one of which we determined the crystal structure of its complex with the SARS-CoV-2 M^pro^. Using this approach we visualized SARS-CoV-2 active M^pro^ within nasopharyngeal epithelial cells of a patient with active COVID-19 infection. The results of our work provide a structural framework for the design of inhibitors as antiviral agents or diagnostic tests.

## Introduction

In December 2019, a severe respiratory disease of unknown origin emerged in Wuhan, Hubei province, China.^1,2^ Symptoms of the first patients were flu-like and included fever, cough and myalgia, but with a tendency to develop a potentially fatal dyspnea and acute respiratory distress syndrome.^2^ Genetic analysis confirmed a betacoronavirus as the causing agent. The virus was initially named 2019 novel coronavirus (2019-nCoV),^1-3^ but shortly thereafter, it was renamed to SARS-CoV-2.^4^ By May 13, 2020, the WHO had registered >4.4 million cumulative cases of coronavirus disease 2019 (COVID-19), with >296.000 deaths.^5^

Currently, there is no approved vaccine or treatment for COVID-19. Efforts are being made to characterize molecular targets, pivotal for the development of anti-coronaviral therapies.^6^ The main protease (M^pro^, also known as 3CL^pro^), is one of the coronavirus non-structural proteins (Nsp5) designated as a potential target for drug development.^7,8^ M^pro^ cleaves the viral polyproteins, generating twelve non-structural proteins (Nsp4-Nsp16), including the RNA-dependent RNA polymerase (RdRp, Nsp12) and the helicase (Nsp13). Inhibition of M^pro^ would prevent the virus from replication and therefore constitutes one of the potential anti-coronaviral strategies.^7-9^

Due to the close phylogenetic relationship between SARS-CoV-2 and SARS-CoV,^3,10,11^ their main proteases share many structural and functional features. From the perspective of the design and synthesis of new M^pro^ inhibitors, a key feature of both the enzymes is their ability to cleave the peptide bond following Gln. The SARS-CoV M^pro^ cleaves polyproteins within the Leu-Gln↓(Ser, Ala, Gly) sequence (↓ indicates the cleavage site), which appears to be a conserved pattern of this protease.^7,9,12^ The specificity for peptide bond hydrolysis after Gln residues is also observed for main proteases of other coronaviruses^13,14^ but is unknown for human enzymes. This observation, along with further studies on the M^pro^, can potentially lead to new broad-spectrum anti-coronaviral inhibitors with minimum side effects.^15^

In the present study, we applied the HyCoSuL (Hybrid Combinatorial Substrate Library) approach to determine the full substrate specificity profile of SARS-CoV and SARS-CoV-2 M^pro^s. The use of natural and a large number of unnatural amino acids with diverse chemical structures allowed an in-depth characterization of the residue preference of the binding pockets within the substrate-binding site of the proteases. The results from library screening enabled us to design and synthesize ACC-labeled substrates with improved catalytic efficiency in comparison to a substrate containing only natural amino acids. Moreover, results from our studies clearly indicate that SARS-CoV M^pro^ and SARS-CoV-2 M^pro^ exhibit highly overlapping substrate specificity. We have used this knowledge to design activity-based probes (ABPs) specific for the SARS-CoV-2 Mpro as well as a peptidomimetic inhibitor. Further, we present a crystal structure of the SARS-CoV-2 M^pro^ in complex with the ABP. Finally, using an ABP, we were able to visualize active SARS-CoV-2 M^pro^ within nasopharyngeal epithelial cells of the patient with active COVID-19 infection. These data provide a useful basis for the design of chemical compounds for effective diagnosis and therapy of COVID-19.

## Results and Discussion

### Substrate specificity of SARS-CoV M^pro^ and SARS-CoV-2 M^pro^

To determine the SARS-CoV M^pro^ and SARS-CoV-2 M^pro^ substrate preferences, we applied a hybrid combinatorial substrate library (HyCoSuL) approach. The library consists of three sublibraries, each of them comprising a fluorescent tag – ACC (7-amino-4-carbamoylmethylcoumarin) –, two fixed positions and two varied positions containing an equimolar mixture of 19 amino acids (Mix) (P2 sublibrary: Ac-Mix-Mix-X-Gln-ACC, P3 sublibrary: Ac-Mix-X-Mix-Gln-ACC, P4 sublibrary: Ac-X-Mix-Mix-Gln-ACC, X =19 natural and over 100 unnatural amino acids, **Figure 1**). We incorporated glutamine at the P1 position, because the available crystal structures of SARS-CoV M^pro^ revealed that only glutamine (and, at only one cleavage site, histidine) can occupy the S1 pocket of this enzyme.^12,16^ The imidazole of His163, located at the very bottom of the S1 pocket, is suitably positioned to interact with the Gln side chain. The Gln is also involved in two other interactions, i.e. with the main chain of F140 and the side chain of Glu166. The library screen revealed that SARS-CoV and SARS-CoV-2 M^pro^ display very similar substrate specificity, however SARS-CoV-2 M^pro^ possesses broader substrate preferences at the P2 position (**Figure 2**). The most preferred amino acid at the P2 position is leucine in case of both proteases. SARS-CoV M^pro^ exhibits lower activity toward other tested amino acids at this position (<30%). Leu selectivity for SARS-CoV M^pro^ is in high agreement with a previous report by Zhu et al..^17^ However, we have noticed some discrepancies in the level of selectivity for other natural amino acids. This is the result of differences in the substrate specificity profiling approach, where we are using a combinatorial mixture of natural amino acids, while Zhu et al. were using a defined peptide sequence. Nevertheless, in both approaches, preferences for the same natural amino acids was observed. The S2 pocket of both investigated enzymes can accommodate other hydrophobic residues, such as 2-Abz, Phe(4-NO_2_), 3-Abz, β-Ala, Dht, hLeu, Met, and Ile (amino acid structures are presented in **Table S1**, SI). At the P3 position, both enzymes prefer hydrophobic D and L amino acids and also positively charged residues; the best are: Tle, D-Phe, D-Tyr, Orn, hArg, Dab, Dht, Lys, D-Phg, D-Trp, Arg, and Met(O)_2_. At the P4 position, SARS-CoV and SARS-CoV-2 M^pro^ possess broad substrate specificity. The most preferred are small aliphatic residues such as Abu, Val, Ala, and Tle, but other hydrophobic amino acids are also accepted. These findings can be partly explained by the available crystal structures of SARS-CoV M^pro^ in complex with inhibitors.^8,12,16^ The hydrophobic S2 subsite of SARS-CoV M^pro^ is more flexible compared to alphacoronavirus M^pro^s, which explains less stringent specificity.^15^ The S2 pocket can form hydrophobic interactions with P2 residues that are not only limited to leucine. The S3 pocket of SARS M^pro^ is not well defined which is also reflected in our P3 substrate specificity profile. The S4 pocket can be occupied by small residues due to the crowded cavity formed by P168, L167 at the bottom and T190, A191 at the top wall.

**Figure 1.**
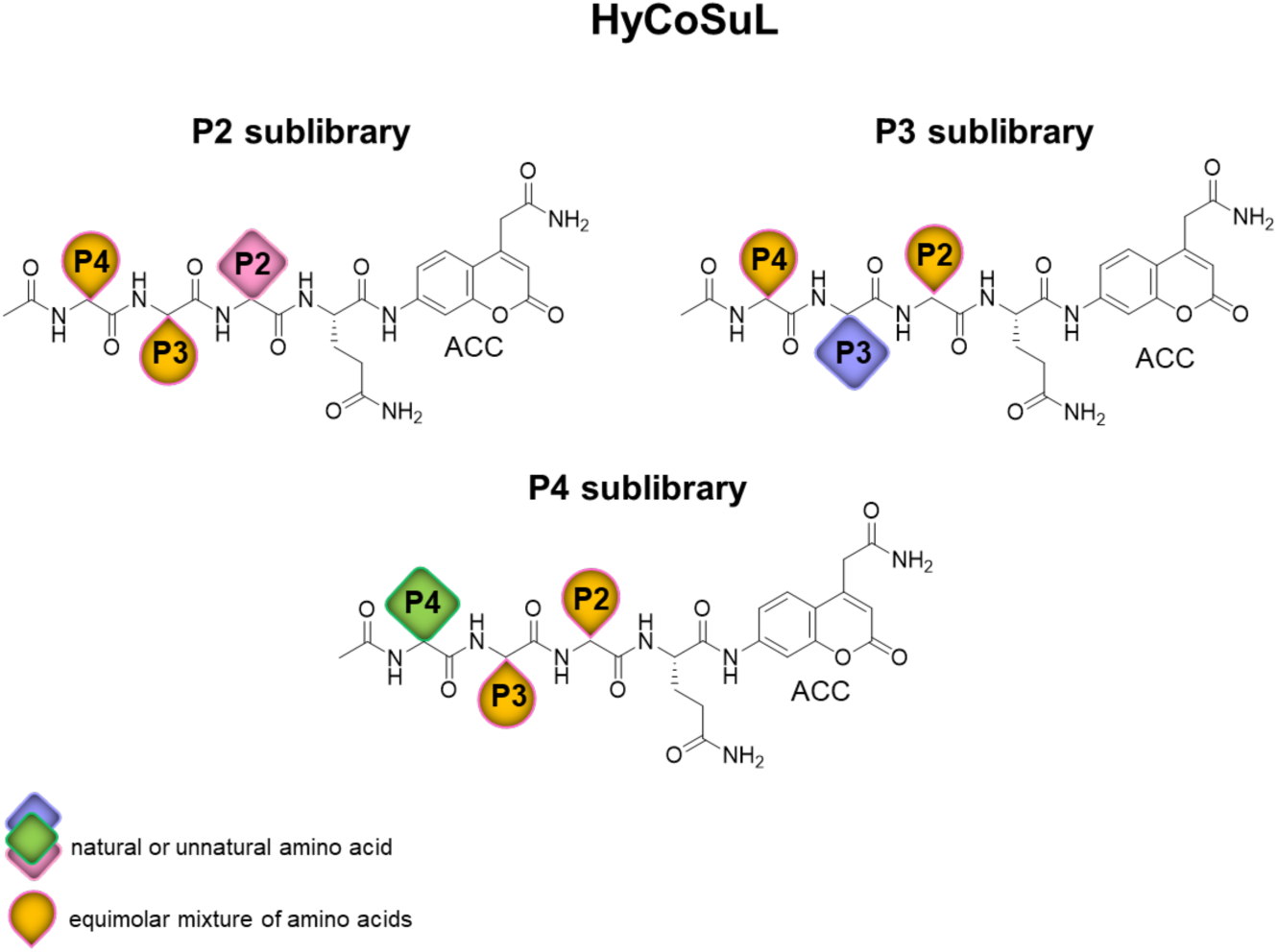
Structure of HyCoSuL library designed for P1-Gln-specific endopeptidases.

**Figure 2.**
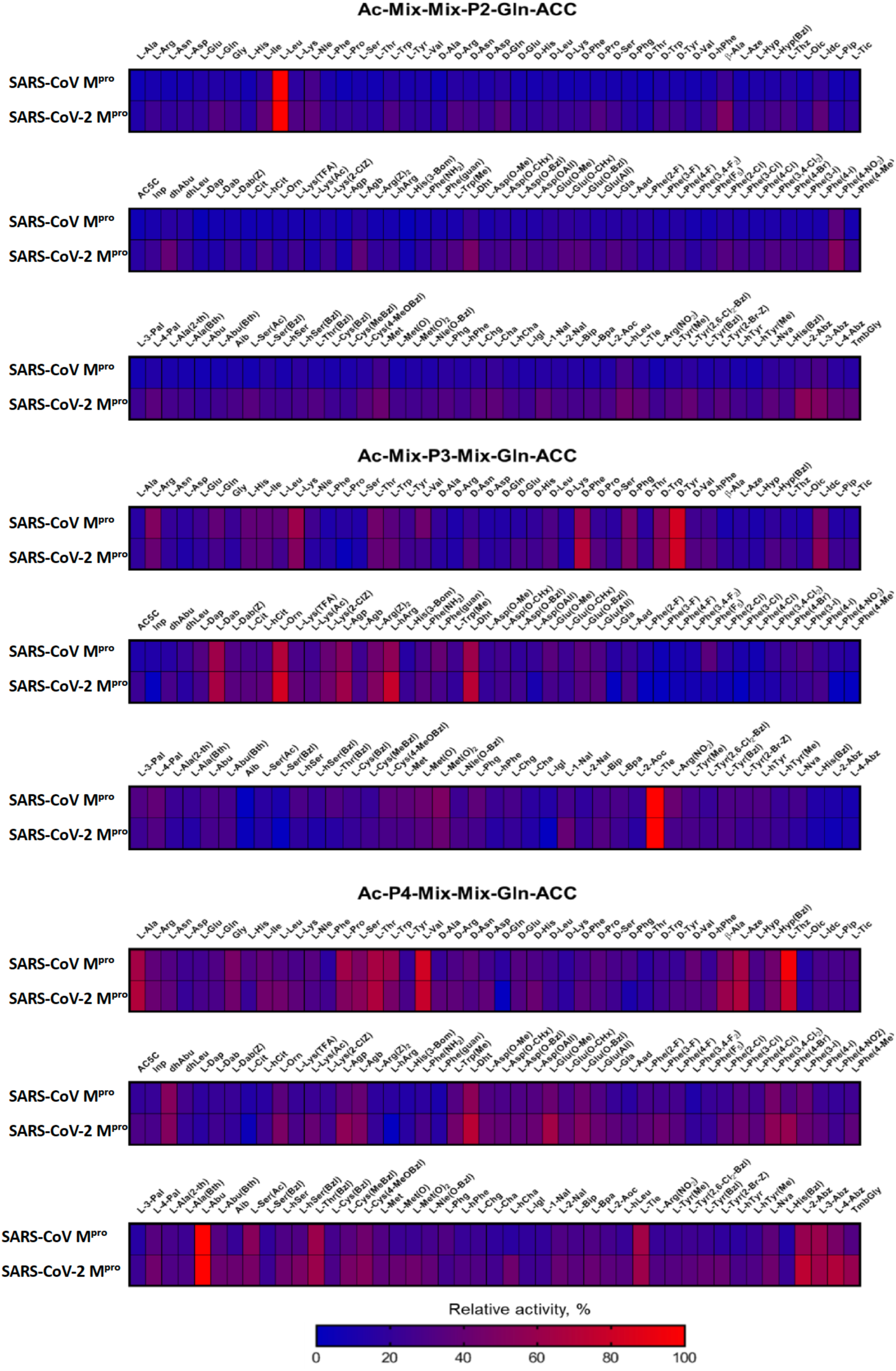
Substrate specificity profiles of SARS-CoV M^pro^ and SARS-CoV-2 M^pro^ presented as heat maps.

To validate the results from library screening, we designed and synthesized ACC-labeled substrates containing the most preferred amino acids in each position. Then, we measured the rate of substrate hydrolysis relevant to each protease (**Figure 3**). The data clearly demonstrate that SARS-CoV M^pro^ and SARS-CoV-2 M^pro^ exhibit the same activity toward tested substrates. The results are consistent with the HyCoSuL screening data. The most preferred substrate, Ac-Abu-Tle-Leu-Gln-ACC, is composed of the best amino acids in each position **(Table 1)**. Kinetic parameters were determined for the two best substrates (Ac-Abu-Tle-Leu-Gln-ACC, Ac-Thz-Tle-Leu-Gln-ACC) and one containing the best recognized natural amino acids (Ac-Val-Lys-Leu-Gln-ACC) (**Table 2**) toward SARS-CoV-2 M^pro^. Due to substrate precipitation because of high concentration needed in the assay, kinetic parameters toward SARS-CoV M^pro^ could not be determined. Analysis of kinetic parameters revealed that these three substrates differ in the k_cat_ value, while K_M_ values are comparable.

**Table 1.**
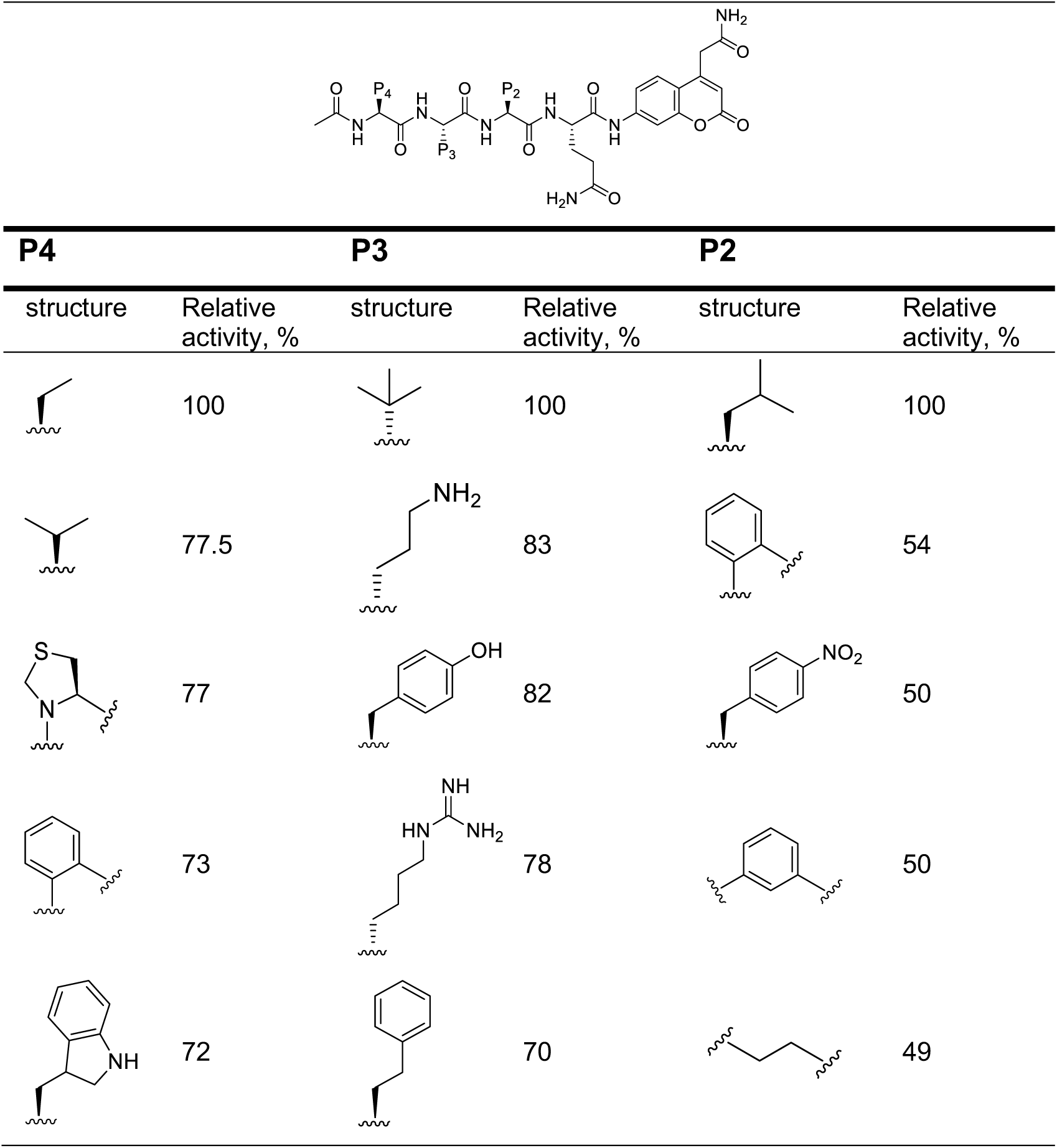
Structures of the most recognizable amino acids at each tested position by SARS-CoV-2 M^pro^.

**Table 2.**
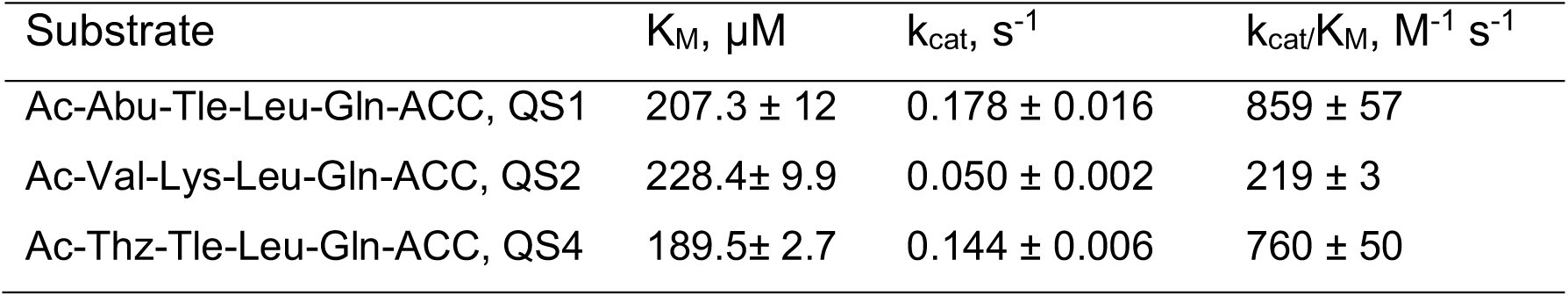
Kinetic parameters of selected substrates for SARS-CoV-2 M^pro^.

| Substrate | K<sub>M</sub>, μM | k<sub>cat</sub>, s<sup>-1</sup> | k<sub>cat</sub>/K<sub>M</sub>, M<sup>-1</sup> s<sup>-1</sup> |
| --- | --- | --- | --- |
| Ac-Abu-Tle-Leu-Gln-ACC, QS1 | 207.3 ± 12 | 0.178 ± 0.016 | 859 ± 57 |
| Ac-Val-Lys-Leu-Gln-ACC, QS2 | 228.4 ± 9.9 | 0.050 ± 0.002 | 219 ± 3 |
| Ac-Thz-Tle-Leu-Gln-ACC, QS4 | 189.5 ± 2.7 | 0.144 ± 0.006 | 760 ± 50 |

**Figure 3.**
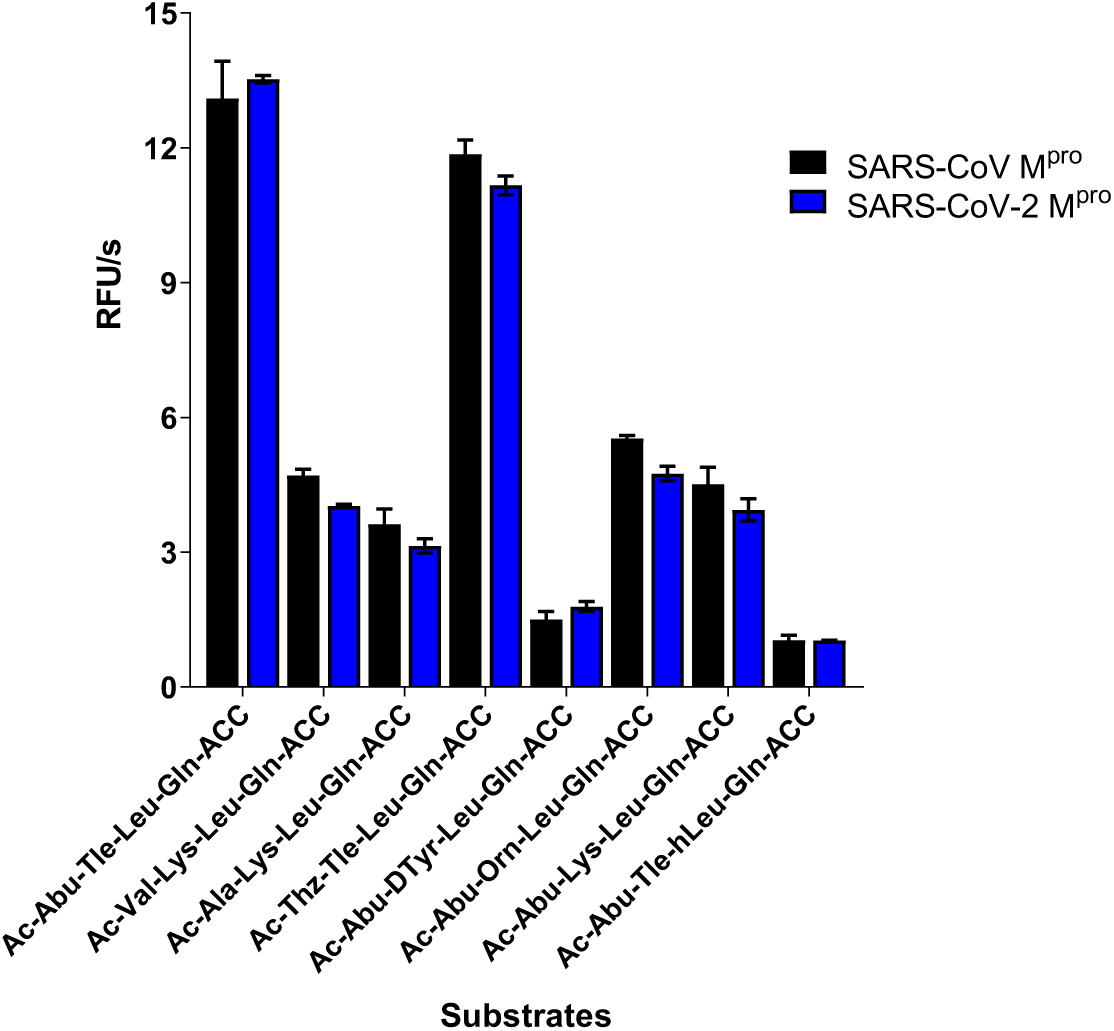
The rate of substrate hydrolysis by SARS-CoV M^pro^ and SARS-CoV-2 M^pro^ ([S]=5 µM, [E]=0.3 µM).

### Inhibitors and activity-based probes of SARS-CoV-2 M^pro^

In the next step, the best substrate QS1 was converted to an inhibitor and activity-based probes. The inhibitor contained the acetylated peptide sequence and vinyl sulfone as an irreversible reactive group (Ac-Abu-Tle-Leu-Gln-VS, Ac-QS1-VS, **Figure 4A**). Probes included a N-terminal biotin tag or Cyanine 5 dye, polyethylene glycol (PEG(4)) as a linker, the best peptide sequence, and vinyl sulfone (**Figure 4A**). To evaluate the sensitivity of designed probes, we performed SDS-PAGE analysis followed by protein transfer onto membranes and ABP visualization. We observed SARS-CoV-2 M^pro^ (100 nM) labelling by Cy5-QS1-VS at a concentration of 100 nM and by B-QS1-VS at 200 nM (**Figure 4B**) which reflected the results of the k_obs_/I analysis (**Table 3**). To determine probe selectivity, we performed cell lysate assays. A HeLa lysate was incubated with different probe concentrations (50, 100, and 200 nM) (**Figure 4C**). The cell lysate experiment confirmed probe selectivity. To verify that unknown bands (about 30 kDa and between 49 and 62 kDa) were due to unspecific Cy5 labelling, we incubated the cell lysate with inhibitor (Ac-QS1-VS) for 30 min at 37°C and then with Cy5-QS1-VS for 15 min at 37°C (lanes 8-10 on the membrane, **Figure 4C**). The same protein bands were observed on the membrane when cell lysates were incubated with and without Ac-QS1-VS, which confirmed unspecific protein labelling by the Cy5 dye.

**Table 3.** Inhibition rate constants of SARS-CoV-2 M^pro^ for inhibitor and activity-based probes.

| Compound | $k_{obs}/I$, M<sup>-1</sup>s<sup>-1</sup> |
| --- | --- |
| Ac-Abu-Tle-Leu-Gln-VS, Ac-QS1-VS | 730 ± 17 |
| Biotin-PEG(4)-Abu-Tle-Leu-Gln-VS, B-QS1-VS | 200 ± 11 |
| Cy5-PEG(4)-Abu-Tle-Leu-Gln-VS, Cy5-QS1-VS | 591 ± 45 |

**Figure 4.**
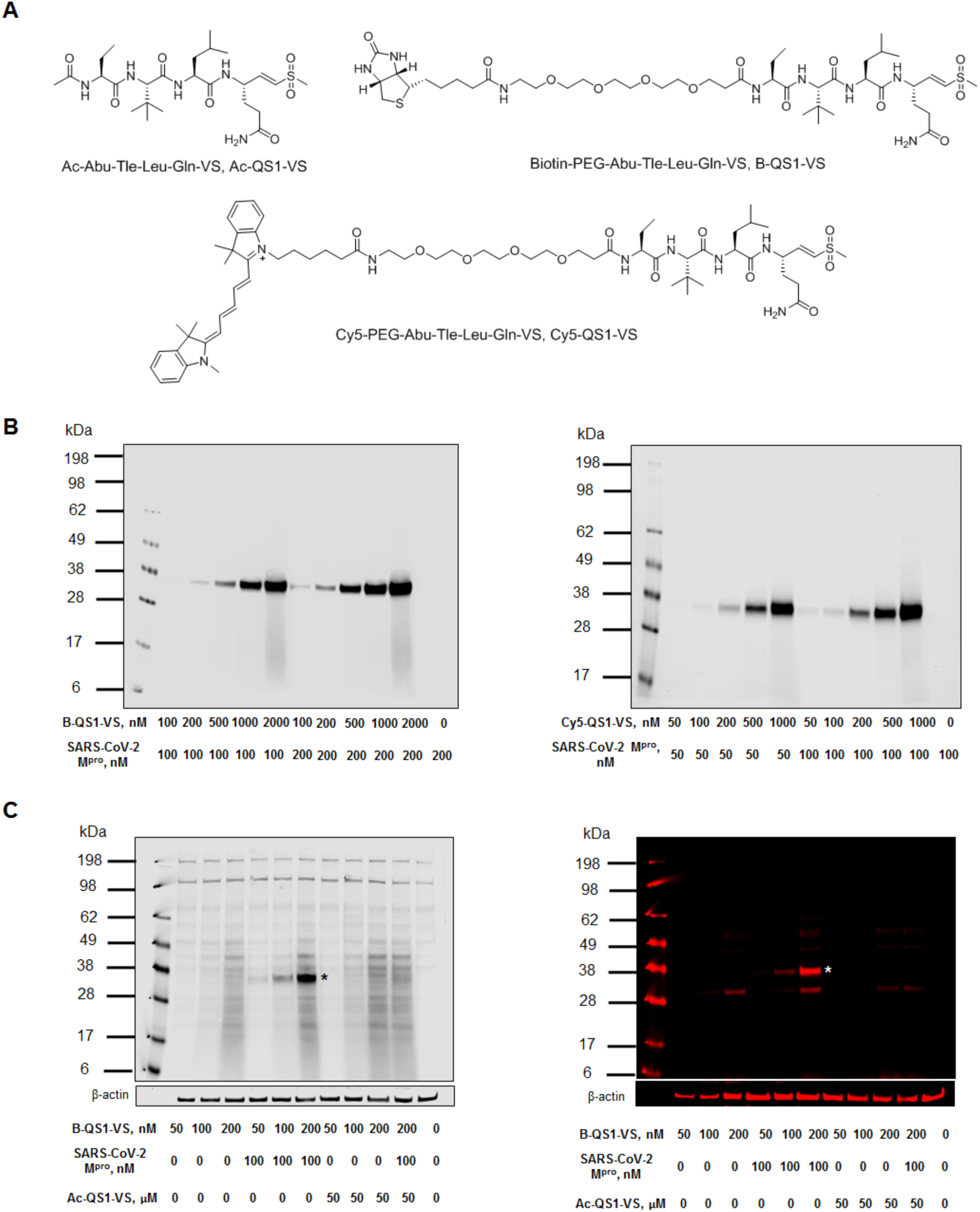
SARS-CoV-2 M^pro^ detection by activity-based probes. (A) Structure of inhibitor and activity-based probes. (B) SARS-CoV-2 M^pro^ labelling by probes (B-QS1-VS and Cy5-QS1-VS). (C) SARS-CoV-2 M^pro^ probe selectivity in HeLa lysate (asterisk shows SARS-CoV-2 M^pro^ band, which was added to the cell lysate). The cell lysate was incubated with or without the inhibitor Ac-QS1-VS for 30 min at 37°C; next, the different probe concentrations were added and the samples were incubated for 15 min at 37°C. The last lane on the membranes is HeLa lysate only.

### Crystal structure of SARS-CoV-2 M^pro^ with the activity-based probe, Biotin-PEG(4)-Abu-Tle-Leu-Gln-VS (B-QS1-VS)

To visualize the steric details of the interactions between the M^pro^ and the activity-based probe, Biotin-PEG(4)-Abu-Tle-Leu-Gln-VS (B-QS1-VS), we determined the X-ray crystal structure of the complex between the two components. The probe was cocrystallized with the recombinant and highly purified SARS-CoV-2 M^pro^. Crystals diffracted to 1.7 Å resolution and were of space group P6_1_22, with one ABP-M^pro^ monomer per asymmetric unit (see Table S2 for crystallographic details). The structure was refined to reasonable R factors and good geometry. Surprisingly, all atoms of the activity-based probe (ABP) were clearly seen in the 2F_o_-F_c_ electron density at a contouring level of 0.5σ. The reason for this is that the flexible tail of the ABP, the PEG(4) chain and the terminal biotin label, are in contact with a neighboring M^pro^ dimer in the crystal lattice (Fig. S1A). This is of course of little relevance for the situation in solution; hence we discuss only the interaction of the P1-P4 residues with the parent M^pro^ molecule (Fig. 5) here. The oxygen atoms of the vinylsulfone group point towards the oxyanion hole of the protease and accept hydrogen bonds (2.81 Å and 3.26 Å) from the main-chain amide groups of Gly143 and Cys145, respectively. Also, the side-chain amide-nitrogen of Asn142 donates a 2.90-Å H-bond to one of these oxygens. The methyl group attached to the sulfone makes hydrophobic contacts with the Cγ2 atom of Thr25 and the side-chain of Leu27, within the S1’ subsite. The catalytic cysteine residue is covalently linked to the Cβ atom of the vinyl group, at a distance of 1.83 Å.

**Figure 5.**
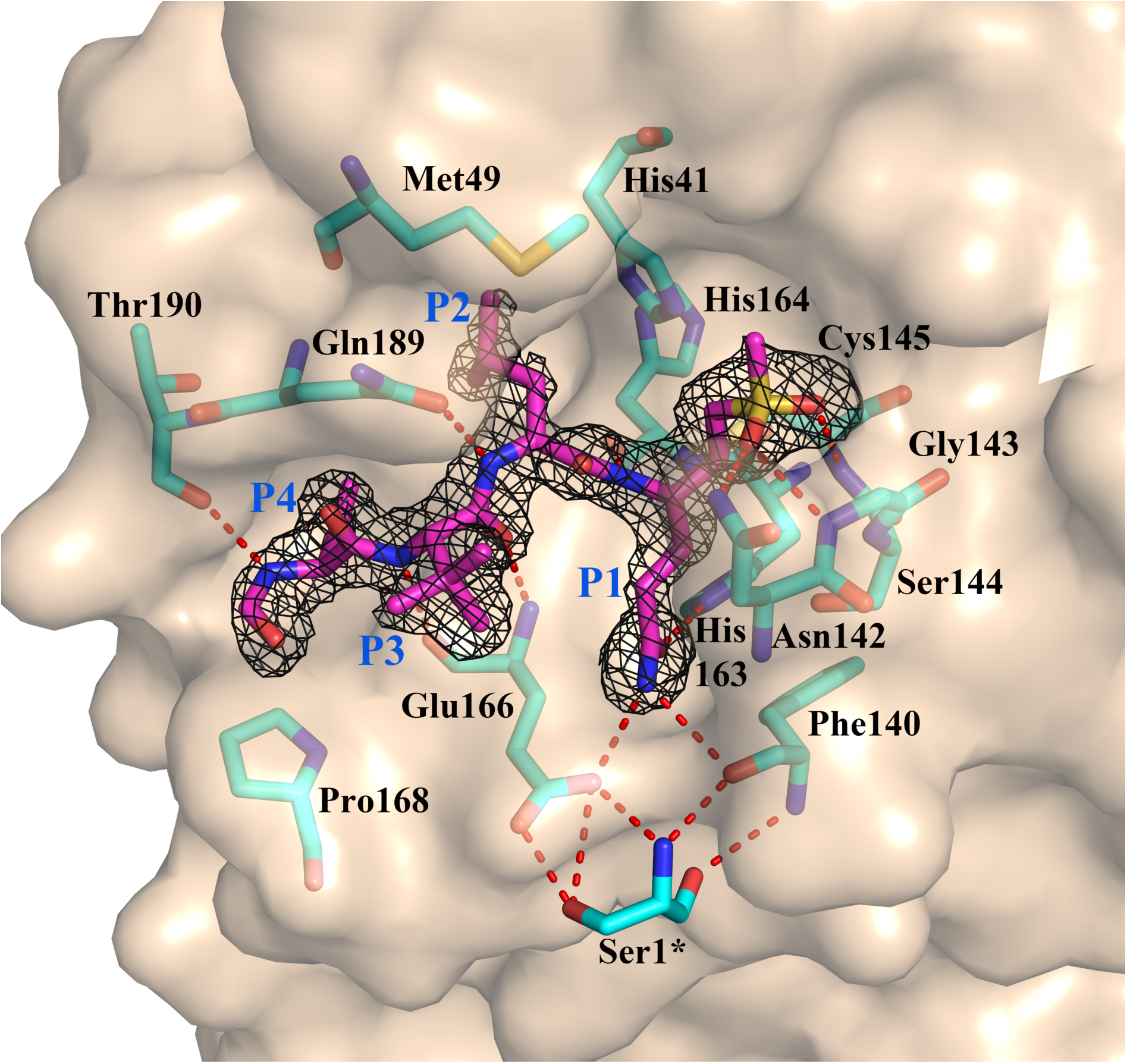
Three-dimensional structure of the P1’ - P4 residues of the activity-based probe (ABP) biotin - PEG(4) - Abu - Tle - Leu - Gln - VS (B-QS1-VS) in the substrate-binding site of the SARS-CoV-2 M^pro^. The F_o_-F_c_ electron density for residues P1’ - P4 of the ABP is shown at a contouring level of 2.5σ. All parts of the ABP beyond the P4 residue would be outside the parent M^pro^ dimer and have been omitted from this figure. However, the rest of the ABP interacts with a neighboring M^pro^ dimer in the crystal (cf. Fig. S1A) and is seen in the electron density maps.

The P1-glutamine side-chain makes the expected interaction with His163 (2.72 Å, through the Oε1 atom) and the Phe140 main-chain oxygen (3.22 Å, through Nε2) and Glu166 Oε1 (3.25 Å). The Nε2 of the P1-Gln thus donates a three-center (bifurcated) hydrogen bond to these two acceptors. This is also reflected in inhibitors that carry a γ-lactam as Gln surrogate in the P1 position, such as compound **13b**.^7^ The Leu residue in the S2 pocket makes the canonical interactions previously observed^17^, with residues Met49, Met165, His41, and the C*α* atom of Arg188. Also, the main-chain amide of the P2-Leu donates a 2.98-Å H-bond to the side-chain Oε1 of Gln189.

There is no well-defined pocket for the P3 moiety, which is therefore mostly solvent-exposed. There may be some weak interaction of the Tle side-chain with the hydrophobic portion of the neighboring Glu166 side-chain. The polar main-chain atoms of the P3 residue form hydrogen-bonds with the protein main chain at Glu166. The aminobutyric acid in P4 makes weak hydrophobic interactions with Leu167 and Gln189, and its main-chain NH group donates a H-bond to Thr190 O.

We have previously noticed that compared to the SARS-CoV M^pro^, Thr285 has been replaced by Ala in SARS-CoV M^pro^, and the neighboring Ile286 by Leu.^7^ In the SARS-CoV M^pro^, the Thr285 makes a hydrogen bond with its symmetry-mate across the two-fold axis creating the M^pro^ dimer. The loss of this H-bond in the SARS-CoV-2 M^pro^ enables the monomers of the dimer to approach each other more closely. Interestingly, in the crystal structure presented here, the space between the two protomers generated by the mutation is filled by a chloride ion adopted from the crystallization buffer (Fig. S1B). This might be taken as a hint for this region around residues 284 - 286 being a hotspot for mutations, due to non-ideal packing of the two protomers of the dimer at this point.

### SARS-CoV-2-Mpro detection and imaging in patient samples

Fluorescent-tagged ABPs currently represent the classic standard in terms of application for labelling of biological samples and have been successfully used for visualization of many proteases in the past^18-20^. We wanted to see if the Cy5-QS1-VS developed by us can be used for detection of SARS-CoV-2 M^pro^ in human samples. Thus, we recruited one patient with mild symptoms of COVID-19, who was positive for SARS-CoV-2 RNA in two independent quantitative RT-PCR assays valid for diagnostic purposes. We incubated the probe at 2 µM final concentration with cells collected from nasopharyngeal swabs of the patient un the first and fifth day after diagnosis and subjected them to confocal laser scanning microscopy. In parallel, we carried out the same experiment with a healthy donor (COVID-19-negative control). These serial measurements revealed that 10-15% of the cells were positive for staining with Cy5-QS1-VS probe in the COVID-19 positive patient (Figure 6). A particularly strong signal from SARS-CoV-2 M^pro^ was observed on the 5^th^ day after diagnosis suggesting a very advanced stage of the infection. No signal from SARS-CoV-2 M^pro^ labelling was observed in the sample from the healthy donor (Figure 6). Thus, we were able to show for the first time that human cells collected *ex vivo* contain active SARS-CoV-2 M^pro^ during SARS-CoV-2 infection.

**Figure 6.**
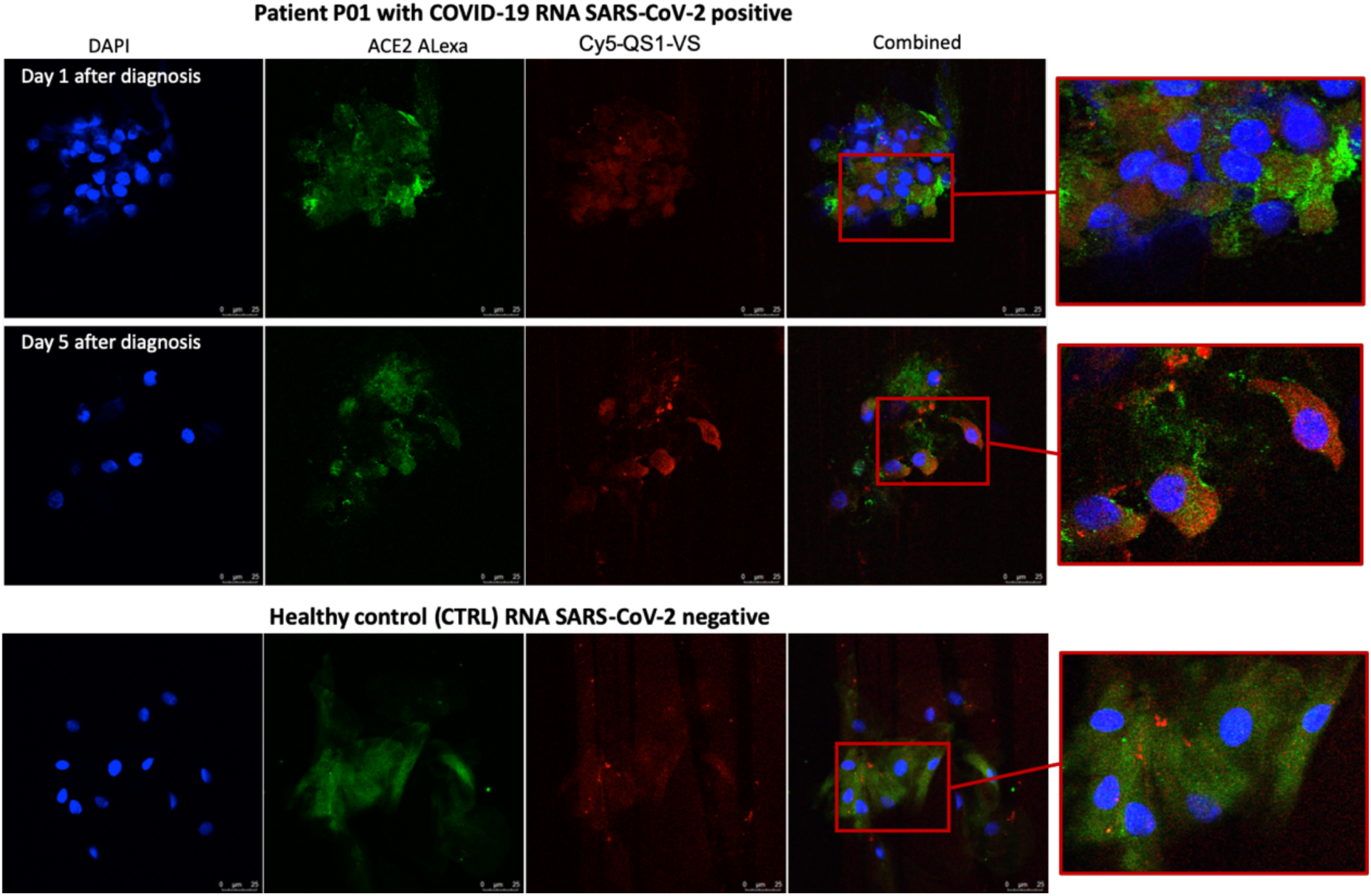
SARS-CoV-2 M^pro^ detection by activity-based probe in nasopharyngeal epithelial cells from COVID-19 patients positive for SARS-CoV-2 RNA (P01). Confocal microscopy of the epithelial cells of nasopharyngeal swabs co-stained with the Cy5-QS1-VS SARS-CoV-2 M^pro^ probe, AF488 anti-ACE2 antibodies, and DAPI in patient P01 at days 1 and 5 after diagnosis as well as in a healthy control.

## Discussion

SARS-CoV-2 first observed in Wuhan, China, caused a global pandemic leading to COVID-19 disease. The lack of a vaccine and approved medications for direct treatment of the disease has led to strenuous efforts to find therapies to stop the pandemic. One of the promising therapeutic targets are two viral proteases - SARS-CoV M^pro^ and SARS-CoV-2-PL^pro^. The first of these, SARS-CoV-2-M^pro^, is used by the virus for protein maturation and its structure has already been described recently^9^. Moreover, the results of retargeting about 10,000 drugs, drugs candidates in clinical trials and other bioactive compounds resulted in selection of several candidates as potential inhibitors of this enzyme^21^. In our research, we decided to thoroughly examine SARS-CoV M^pro^ to find the optimal chemical tools in the form of substrates, inhibitors and activity-based probes. First, we have obtained a targeted library of fluorogenic substrates (HyCoSuL) towards this protease and determined the substrate specificity at the P4-P2 positions. We have directly compared the substrate specificity of the main protease with the same protease from previous SARS, which caused an epidemy in 2003. Our data clearly demonstrate that these enzymes have very similar preferences for natural and unnatural amino acids in the P4-P2 positions. They tolerate many different amino acids at P4 and P3, and have a strong preference for Leu at P2. This information is certainly crucial for the aspect of drug retargeting, but also the use of information obtained in previous years for SARS-CoV M^pro^ to be used for current research. In the next step, we created potent inhibitors and activity-based probes, whose sequences were based on the HyCoSuL screening results. In the case of B-QS1-VS, we also obtained a crystal structure from SARS-CoV-2 M^pro^, which accurately shows the binding mechanism in the P4-P1’ pockets. This knowledge certainly complements the information already obtained for other inhibitor molecules published for this enzyme. In turn, we used the fluorescent activity-based probe to visualize the SARS-CoV-2 M^pro^ activity in patient samples with COVID-19, thus confirming that it is an excellent tool that can be used to detect this enzyme, and also be used as a diagnostic tool.

In conclusion, our research allowed the exact characterization of SARS-CoV-2 M^pro^ in terms of both amino-acid preferences as well as design of targeted inhibitors and activity-based probes. The reagents described here can be used to optimize structures that lead to anti-COVID-19 drugs, as well as further drug retargeting.

## Experimental Section

### Reagents

The reagents used for solid-phase peptide synthesis were as follows: Rink Amide (RA) resin (particle size 100-200 mesh, loading 0.74 mmol/g), all Fmoc-amino acids, *O*-benzotriazole-*N,N,N’,N’*-tetramethyl-uronium-hexafluoro-phosphate (HBTU), 2-(1-H-7-azabenzotriazol-1-yl)-1,1,3,3-tetramethyluranium hexafluorophosphate (HATU), piperidine, diisopropylcarbodiimide (DICI), and trifluoroacetic acid (TFA), purchased from Iris Biotech GmbH (Marktredwitz, Germany); anhydrous *N*-hydroxybenzotriazole (HOBt) from Creosauls, Louisville, KY, USA; 2,4,6-collidine (2,4,6-trimethylpyridine), HPLC-grade acetonitrile, triisopropylsilane (TIPS) from Sigma-Aldrich (Poznan, Poland); and *N,N*-diisopropylethylamie (DIPEA) from VWR International (Gdansk, Poland). *N,N*-dimethylformamide (DMF), dichloromethane (DCM), methanol (MeOH), diethyl ether (Et_2_O), acetic acid (AcOH), and phosphorus pentoxide (P_2_O_5_), obtained from Avantor (Gliwice, Poland). Designed substrates were purified by HPLC on a Waters M600 solvent delivery module with a Waters M2489 detector system using a semipreparative Wide Pore C8 Discovery column. The solvent composition was as follows: phase A (water/0.1% TFA) and phase B (acetonitrile/0.1% TFA). The purity of each compound was confirmed with an analytical HPLC system using a Jupiter 10 µm C4 300 Å column (250 x 4.6 mm). The solvent composition was as follows: phase A (water/0.1% TFA) and phase B (acetonitrile/0.1% TFA); gradient, from 5% B to 95% B over a period of 15 min. The purity of all compounds was ≥95%. The molecular weight of each substrate was confirmed by high-resolution mass spectrometry using a Waters LCT premier XE with electrospray ionization (ESI) and a time-of-flight (TOF) module.

### Enzyme preparation

Gene cloning and recombinant production of the SARS-CoV and SARS-CoV-2 M^pro^ are described elsewhere.^9,17^

### Combinatorial library synthesis

#### Synthesis of H_2_N-ACC-resin

ACC synthesis was carried out according to Maly et al.^22^ To a glass reaction vessel, 1 eq (9.62 mmol, 13 g) of Rink AM resin was added and stirred gently once per 10 min in DCM for 1 h, then filtered and washed 3 times with DMF. Fmoc-group deprotection was performed using 20% piperidine in DMF (three cycles: 5, 5, and 25 min), filtered and washed with DMF each time (six times). Next, 2.5 eq of Fmoc-ACC-OH (24.05 mmol, 10.64 g) was preactivated with 2.5 eq HOBt monohydrate (24.05 mmol, 3.61 g) and 2.5 eq DICI (24.05 mmol, 3.75 mL) in DMF and the slurry was added to the resin. The reaction was shaked gently for 24 hours at room temperature. After this time, the resin was washed four times with DMF and the reaction was repeated using 1.5 eq of above reagents in order to improve the yield of ACC coupling to the resin. After 24 hours, the resin was washed with DMF and the Fmoc protecting group was removed using 20% piperidine in DMF (5, 5, and 25 min), filtered and washed with DMF (six times).

#### Synthesis of H_2_N-Gln(Trt)-ACC-resin

2.5 eq Fmoc-Gln(Trt)-OH (24.05 mmol, 14.69 g) with 2.5 eq HATU (24.05 mmol, 9.15 g), 2.5 eq collidine (24.05 mmol, 3.18 mL) in DMF were activated for 2 min and added to filter cannula with 1 eq (9.62 mmol) H_2_N-ACC-resin and the reaction was carried out for 24 h. Next, the resin was washed four times with DMF and the same reaction was performed again using 1.5 eq of above reagents. After four DMF washes, the Fmoc protecting group was removed using 20% piperidine in DMF (5, 5, and 25 min). Subsequently, the resin was washed with DCM (3 times) and MeOH (3 times) and dried over P_2_O_5_. The synthesis of P2, P3, and P4 sublibraries is exemplified in detail with the P2 sublibrary. The P2 library consisted of 137 compounds where all of the natural amino acids (omitting cysteine) and a pool of unnatural amino acids were used at a defined position (in this case, the P2 position) and an isokinetic mixture of 19 amino acids (without cysteine; plus norleucine mimicking methionine) was coupled in the remaining positions (in case of the P2 sublibrary, positions P3 and P4 were occupied by an isokinetic mixture). Equivalent ratios of amino acids in the isokinetic mixture were defined based on their reported coupling rates. A fivefold excess (over the resin load) of the mixture was used. For fixed positions, 2.5 eq of single amino acid was used. All reactions were performed with the use of the coupling reagents DICI and HOBt. For P2 coupling, the synthesis of the library was performed using a MultiChem 48-well synthesis apparatus (FlexChem from SciGene, Sunnyvale, CA, USA). To each well of the reaction apparatus, 1 eq of dry H_2_N-Gln(Trt)-ACC-resin (0.059 mmol, 80 mg) was added and stirred gently for 30 minutes in DCM, and then washed four times with DMF. In separate Eppendorf tubes, 2.5 eq (0.15 mmol) Fmoc-P2-OH was preactivated with 2.5 eq HOBt (0.15 mmol, 22.5 mg) and 2.5 eq DICI (0.15 mmol, 23.55 μL) in DMF. Next, preactivated amino acids were added to wells of the apparatus containing H_2_N-Gln(Trt)-ACC-resin, followed by 3 h of agitation at room temperature. Then, the reaction mixture was filtered, washed with DMF (4 times), and the ninhydrin test was carried out in order to confirm P2-amino acid coupling. Subsequently, Fmoc protecting groups were removed with the use of 20% piperidine in DMF (5, 5, and 25 min). For P3 and P4 position coupling, an isokinetic mixture for 48 portions was prepared from 18 Fmoc-protected natural amino acids (omitting cysteine; plus norleucine mimicking methionine; 19 amino acids in total). Next, 5 eq of isokinetic mixture, 5 eq HOBt (14.16 mmol, 2.13 g), and 5 eq DICI (14.16 mmol, 2.22 mL) were diluted in DMF and preactivated for 3 min. The activated isokinetic mixture was added to each of 48 wells containing 1 eq of H_2_N-P2-Gln(Trt)-ACC-resin. After 3 h of gentle agitation, the slurry was filtered off and washed with DMF (4 times). A ninhydrin test was carried out and the Fmoc protecting group was removed using 20% piperidine in DMF (5, 5, and 25 min). The same procedure was applied for the remaining compounds. The isokinetic mixture was added to prepare the P4 position in the same manner as for the P3 position. In the last step of the synthesis, *N*-terminal acetylation was performed; to prepare the mixture for 48 compounds, 5 eq of AcOH (14.16 mmol, 807 µL), 5 eq HBTU (14.16 mmol, 5.37 g), and 5 eq DIPEA (14.16 mmol, 2.44 mL) in ∼45 mL of DMF were added to a 50-mL falcon tube. After gentle stirring for 1 min, the mixture (∼800 µL) was added to each well in the reaction apparatus, containing the H_2_N-Mix-Mix-P2-Gln(Trt)-ACC-resin, followed by gentle agitation for 30 min. Next, the resin was washed six times with DMF, three times with DCM, three times with MeOH, and dried over P_2_O_5_. After completing the synthesis, peptides were cleaved from the resin with a mixture of cold TFA:TIPS:H2O (%, v/v/v 95:2.5:2.5; 2 mL/well; 2 hours, shaking once per 15 min). The solution from each well was collected separately and the resin was washed once with a portion of fresh cleavage solution (1 mL), followed by addition of diethyl ether (Et_2_O, 14 mL) into falcons with peptides in solution. After precipitation (30 min at - 20°C), the mixture was centrifuged and washed again with Et_2_O (5 mL). After centrifugation, the supernatant was removed and the remaining white precipitate was dissolved in ACN/H_2_O (v/v, 3/1) and lyophilized. The products were dissolved in DMSO to a final concentration of 10 mM and used without further purification. The synthesis of P3 and P4 sublibraries was performed in the same manner as described above; P3 and P4 sublibraries were synthesized by coupling fixed amino-acid residues to P3 (isokinetic mixture coupled to P2 and P4) and P4 position (isokinetic mixture coupled to P2 and P3).

### Library screening

Hybrid combinatorial substrate library screening was performed using a spectrofluorometer (Molecular Devices Spectramax Gemini XPS) in 384-well plates (Corning). The assay conditions were as follows: 1 µL of substrate and 49 µL of enzyme, which was incubated at 37°C for 10 min in assay buffer (20 mM Tris, 150 mM NaCl, 1 mM EDTA, 1 mM DTT, pH 7.3). The final substrate concentration was 100 µM and the final enzyme concentration was 1 µM SARS-CoV and 0.6 µM SARS-CoV-2 M^pro^, respectively. The release of ACC was measured for 45 min (λ_ex_ = 355 nm, λ_em_ = 460 nm) and the linear part of each progress curve was used to determine the substrate hydrolysis rate. Substrate specificity profiles were established by setting the highest value of relative fluorescence unit per second (RFU/s) from each position as 100% and others were adjusted accordingly.

### Individual substrate synthesis

ACC-labeled substrates were synthesized on the solid support according to the solid phase peptide synthesis method described elsewhere.^18^ In brief, Fmoc-ACC-OH (2.5 eq) was attached to a Rink-amide resin using HOBt (2.5 eq) and DICI (2.5 eq) in DMF as coupling reagents. Then, the Fmoc protecting group was removed using 20% piperidine in DMF (three cycles: 5, 5, and 25 min). Fmoc-Gln(Trt)-OH (2.5 eq) was coupled to the H_2_N-ACC-resin using HATU (2.5 eq) and 2,4,6-collidine (2.5 eq) in DMF. After Fmoc group removal, Fmoc-P2-OH (2.5 eq) amino acid was attached (HOBt and DICI (2.5 eq) in DMF). Amino acids in P3 and P4 positions were coupled in the same manner. The free *N*-terminus was acetylated using HBTU, AcOH and DIPEA in DMF (5 eq of each reagent). Then, the resin was washed five times with DMF, three times with DCM and three times with MeOH, and dried over P_2_O_5_. Substrates were removed from the resin with a mixture of TFA/TIPS/H_2_O (% v/v/v, 95:2.5:2.5), precipitated in Et_2_O, purified on HPLC and lyophilized. The purity of each substrate was confirmed using analytical HPLC. Each substrate was dissolved in DMSO at a final concentration of 10 mM and stored at -80°C until use.

### Kinetic analysis of substrates

Substrate screening was carried out in the same manner as the library assay. Substrate concentration was 5 µM, SARS-CoV M^pro^ was 0.3 µM and SARS-CoV-2 M^pro^ was 0.3 µM. Substrate hydrolysis was measured for 30 min using the following wavelengths: λ_ex_ = 355 nm, λ_em_ = 460 nm. The experiment was repeated three times. Results were presented as mean values with standard deviations. Kinetic parameters were assayed in 96-well plates (Corning). Wells contained 80 µL of enzyme in assay buffer (0.074-0.1 µM SARS-CoV-2 M^pro^) and 20 µL of substrate at eight different concentrations ranging from 58.5 µM to 1200 µM. ACC liberation was monitored for 30 min (λ_ex_ = 355 nm, λ_em_ = 460 nm). Each experiment was repeated at least three times. Kinetic parameters were determined using the Michaelis-Menten equation and GraphPad Prism software.

### Activity-based probe and inhibitor synthesis

The synthesis of the biotinylated activity-based probe involved three sequential steps. In the first step, the Biotin-PEG(4)-Abu-Tle-Leu-OH fragment was synthesized using a 2-chlorotrityl chloride resin. Fmoc-Leu-OH (2.5 eq) was dissolved in anhydrous DCM, pre-activated with DIPEA (3 eq) and added to the cartridge with resin (1 eq). After 3h, the mixture was filtered, washed 3 times with DCM and 3 times with DMF, and the Fmoc group was removed using 20% piperidine in DMF. Fmoc-Tle-OH, Fmoc-Abu-OH and Fmoc-PEG(4)-OH were attached to the resin using HATU (2.5 eq) and 2,4,6-collidine (2.5 eq) in DMF as coupling reagents. The biotin tag was coupled to H_2_N-PEG(4)-Abu-Tle-Leu-resin using 2.5 eq HBTU and 2.5 eq DIPEA in a DMF:DMSO mixture (1:1, v/v). After 3 h, the resin was washed 3 times with DMF, 3 times with DCM, and 3 times with MeOH and dried over P_2_O_5_. Finally, the peptide fragment was removed from the resin with a mixture of DCM/TFE/AcOH (v/v/v, 8:1:1). The solution was filtered and concentrated. The obtained crude peptide was dissolved in ACN:H_2_O (v/v, 3:1), lyophilized, and then used without further purification (purity > 95%). In the second step, Fmoc-Gln(Trt)-VS was synthesized in the same manner as described elsewhere^23^. In the third step, the Fmoc group was removed from Fmoc-Gln(Trt)-VS using a mixture of diethylamine:ACN (1:1, v/v). Biotin-PEG(4)-Abu-Tle-Leu-OH was pre-activated with HATU (1.2 eq) and 2,4,6-collidine (3 eq) in DMF and attached to the H_2_N-Gln(Trt)-VS (1 eq). The reaction was agitated for 2 h and the product was purified on HPLC. Finally, Biotin-PEG(4)-Abu-Tle-Leu-Gln(Trt)-VS was treated with a mixture of TFA/DCM/TIPS (% v/v/v/ 70:27:3) to remove the Trt group. After 40 min, solvents were evaporated and the probe was purified on HPLC.

Ac-Abu-Tle-Leu-Gln-VS and Cy5-PEG(4)-Abu-Tle-Leu-Gln-VS were synthesized in the same manner, but Boc-PEG(4)-Abu-Tle-Leu-OH and Ac-Abu-Tle-Leu-OH were synthesized on the 2-chlorotrityl chloride resin instead of Biotin-PEG(4)-Abu-Tle-Leu-OH.

### Determination of inhibition kinetics (k_obs_/I) for inhibitor and activity-based probes

SARS-CoV-2 M^pro^ (75 nM) was preincubated in assay buffer 20 mM Tris, 150 mM NaCl, 1 mM EDTA, 1 mM DTT, pH 7.3 for 10 min at 37°C. Then, the enzyme was added to wells containing seven different concentrations of inhibitor or probe (ranging from 3.9 µM to 20 µM) and 50 μM of substrate (QS1). The measurement was conducted for 30 minutes and repeated at least three times; k_obs_/I was calculated as previously described.^24^

### SARS-CoV-2 M^pro^ labelling

SARS-CoV-2 M^pro^ (50, 100, or 200 nM) was incubated with different probe concentrations (50-2000 nM) in assay buffer (20 mM Tris, 150 mM NaCl, 1 mM EDTA, 1 mM DTT, pH 7.3) for 15 min at 37°C. Then 3x SDS/DTT was added, and the samples were boiled for 5 min at 95°C and resolved on 4-12% Bis-Tris Plus 12-well gels at 30 µL sample/well. Electrophoresis was performed at 200 V for 29 min. Next, the proteins were transferred to a nitrocellulose membrane (0.2 µm, Bio-Rad) for 60 min at 10 V. The membrane was blocked with 2% BSA in Tris-buffered saline with 0.1% (v/v) Tween 20 (TBS-T) for 60 min at RT. The biotinylated activity-based probe was detected with a fluorescent streptavidin Alexa Fluor 647 conjugate (1:10000) in TBS-T with 1% BSA using an Azure Biosystems Sapphire Biomolecular Imager and Azure Spot Analysis Software.

### SARS-CoV-2 M^pro^ labelling in HeLa lysates

HeLa cells were cultured in DMEM supplemented with 10% fetal bovine serum, 2 mM L-glutamine, and antibiotics (100 U/mL penicillin, 100 µg/mL streptomycin) in a humidified 5% CO_2_ atmosphere at 37°C. Approximately 1,200,000 cells were harvested and washed three times with PBS. The cell pellet was lysed in buffer containing 20 mM Tris, 150 mM NaCl, and 5 mM DTT, pH 8.0, using a sonicator. The cell lysate was centrifuged for 10 min, and the supernatant was collected. Twenty microliters of cell lysate was incubated with or without 30 µL of inhibitor Ac-QS1-VS and with or without SARS-CoV-2 M^pro^ (100 nM) for 30 min at 37°C. Next, 50 µL of B-QS1-VS or Cy5-QS1-VS at different concentrations was added to the samples and they were incubated for 15 min at 37°C. Then the samples were combined with 50 µL 3xSDS/DTT, boiled, and run on a gel. Electrophoresis, protein transfer to a nitrocellulose membrane, and probe visualization were conducted in the same manner as described above.

### Crystallization of SARS-CoV-2 M^pro^ in complex with biotin-PEG(4)-Abu-Tle-Leu-Gln-vinylsulfone

The purified SARS-CoV-2 M^pro^ was concentrated to 23 mg/mL, mixed with biotin-PEG(4)-Abu-Tle-Leu-Gln-vinylsulfone at a molar ratio of 1:5, and the mixture was incubated at 4°C overnight. The next day, the mixture was clarified by centrifugation at 12,000 x *g*, 4°C, and the supernatant was set for crystallization screening by using commercially available screening kits (PEGRx™ 1 & 2 (Hampton Research) and Morpheus HT-96 (Molecular Dimensions)). A Gryphon LCP crystallization robot (Art Robbins) was used for setting up the crystallization screens with the sitting-drop vapor-diffusion method at 18°C, where 0.15 μL of protein solution and 0.15 μL of reservoir were mixed to equilibrate against 40 μL reservoir solution.

Crystals were observed under several conditions in the two 96-well plates. The crystals were fished directly from the basic screening plates. The cryo-protectant consisted of mother liquor plus varied concentrations (5% to 20%) of glycerol, and 2 mM of the activity-based probe (ABP). Subsequently, liquid nitrogen was used for flash-cooling the crystals prior to data collection.

### Diffraction data collection and determination of the structure

Several diffraction data sets were collected at 100 K at the P11 beamline of PETRA III (DESY, Hamburg, Germany), using synchrotron radiation of wavelength 1.0332 Å and a Pilatus 6M detector (Dectris). For structure determination, a data set was used that was collected using a crystal fished from the condition No. E7 of Morpheus HT-96 (0.12 M ethylene glycols (0.3 M diethylene glycol, 0.3 M triethylene glycol, 0.3 M tetraethylene glycol, 0.3 M pentaethylene glycol), 0.1 M buffer system 2 (1.0 M sodium HEPES, MOPS (acid), pH 7.5), pH 7.5, 30% precipitant mix 3 (20% v/v glycerol, 10% PEG 4000)).

*XDSapp*,^*25*^ *Pointless*,^*26,27*^ and *Scala*^*26*^ (the latter two from the CCP4 suite^28^) were used for processing and scaling the dataset. The space group was determined as *P*6_1_22 and the resolution limit was set at a Bragg spacing of 1.70 Å. The molecular replacement method was employed for phase determination using the *Molrep* program^28,29^ and the free-enzyme crystal structure of SARS-CoV-2 M^pro^ (PDB entry 6Y2E;^9^) as a the search model. The geometric restraints for the activity-based probe were generated using the *Jligand* programe from the CCP4 suite;^28,30^ the ABP was built into the F_o_-F_c_ density by using the *Coot* software.^31^ Structure refinement was performed with *Refmac5.*^*28,31,32*^ Statistics of diffraction data processing and model refinement are shown in Table S2.

### Patient sample preparation

The study was approved by the Bioethics Committee of the Medical University of Lodz, Poland (# RNN/114/20/KE). The study was conducted in compliance with good clinical practice guidelines and under the principles of the Declaration of Helsinki. The study participants or their parents provided written informed consent. A 13-year old boy, who was positive for SARS-CoV-2 RNA in serial measurements, was included in the study. This subject presented mild symptoms of COVID-19 infection. Two nasopharyngeal swabs were collected twice during routine clinical examination, at day 1 and 5 after COVID-19 diagnosis (positive SARS-CoV-2 RNA test).

### Detection of SARS-CoV-2 M^pro^ in epithelial cells by immunofluorescence

The nasopharyngeal swab was used for smear preparation for confocal laser scanning microscopy. Smears were performed on glass slides covered with polylysine (Thermo Fisher Scientific, MA, USA). These slides were treated with 2 μM probe of M^pro^ Cy5-QS1-VS and incubated for 30 min at 37°C. Slides were fixed in 70% ethanol for 30 min. For further staining overnight at +4°C, incubation with recombinant goat antiACE2 antibody (R&D Systems, MN, USA, 1:20) was performed. The next day, antibodies were aspirated, and cells were washed twice with PBS and labeled with a secondary antibody (Alexa Fluor 488 donkey anti-Goat IgG (H+L) (A11055, 1:200, Life Technologies) in PBS for 1 hour at room temperature. Next, the cells were washed twice with PBS, coverslips were mounted with Vectashield fluorescence mounting medium containing DAPI (Vector Lab. H-1000) and sealed with nail polish. Slides were stored at +4°C until use. Cells were then subjected to confocal microscope analysis using a Leica TCS SP8. DAPI was detected by the DAPI channel (405 nm), the M^pro^ Cy5-QS1-VS probe was detected with the Cy5 filter single photon laser (658 nm), and ACE2/secondary antibodies were read using the FITC filter (single photon laser: 458 nm). All images were acquired in .tiff format using Leica Application Suite X software. Images shown are representative views of cells from two coverslips.

## Ancillary Information

### Supporting Information

Structures of natural and unnatural amino acids, crystal structure data and characterization of all compounds are included in the supplementary materials.

### Author contributions

M. D. and W.R. designed the research; W.R., K.G., L.Z. and M.Z. performed the research and collected data; R.H., X.S. and L.Z. contributed enzymes; L.Z. and R.H. solved and refined the crystal structure; B.P. and W.M. carried out labelling experiments in patient samples and confocal imaging; W.R., L.Z., W.M. R.H. and M.D. analyzed and interpreted the data; W.R., L.Z., R.H. and M.D. wrote the manuscript; and all authors critically revised the manuscript.

## Acknowledgments

The Drag laboratory is supported by the National Science Centre in Poland and the “TEAM/2017-4/32” project, which is conducted within the TEAM programme of the Foundation for Polish Science cofinanced by the European Union under the European Regional Development Fund. Work in the Hilgenfeld laboratory was supported by the SCORE project of the European Union (grant agreement # 101003627) and by the Government of Schleswig-Holstein through its Structure and Excellence Fund, as well as by a close partnership between the Possehl Foundation (Lübeck) and the University of Lübeck. L.Z. is supported by a stipend from the German Center for Infection Research (DZIF). W.R. is a beneficiary of a START scholarship from the Foundation for Polish Science.

## Abbreviations used

Abu: 2-aminobutanoic acid;
2-Abz: 2-(amino)benzoic acid;
3-Abz: 3-(amino)benzoic acid;
ACC: 7-amino-4-carbamoylmethylcoumarin;
Dab: 2,4-diaminobutyric acid;
Dht: dihydrotryptophan;
HyCoSuL: Hybrid Combinatorial Substrate Library;
Orn: ornithine;
RFU: relative fluorescence unit;
D-Phg: D-phenylglycine;
Thz: thiazolidine-4-carboxylic acid;
Tle: tert-leucine;

