## Supporting data for "Substrate specificity profiling of SARS-CoV-2 main protease enables design of activity-based probes for patient-sample imaging"

Content of Supporting Information

1. Figure S1…………………………………………………………………………… S2
2. Supplementary Tables……………………………………..……………………....S3-S12
2. Analysis data of synthesized substrates…………………………..………...……..S13-S23

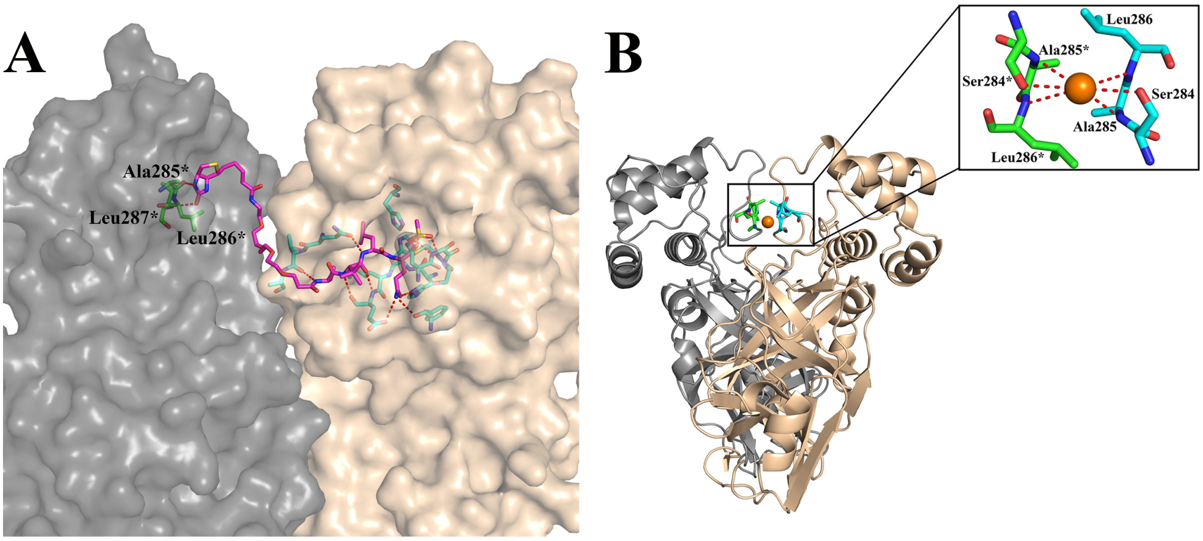


**Figure S1.** Three-dimensional structure of the activity-based probe (ABP) biotin - PEG(4) - Abu - Tle - Leu - Gln - VS (B-QS1-VS) in complex with the SARS-CoV-2 $M^{pro}$. **A**. In the crystal, the ABP links two neighboring $M^{pro}$ dimers. The vinylsulfone warhead and the P1 - P4 residues bind to the substrate-binding site of the parent $M^{pro}$ dimer (wheat), while the PEG(4) unit and the biotin label are in contct with the neighboring $M^{pro}$ dimer (grey). B. At the monomer - monomer interface of the $M^{pro}$, a chloride ion from the crystallization buffer occupies a position on the two-fold axis.

**Table S1.** Structures of fixed natural and unnatural amino acids used in combinatorial library.

| No | Structure + code | No | Structure + code | No | Structure + code |
|---|---|---|---|---|---|
| 1 | 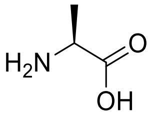 L-Ala | 2 | 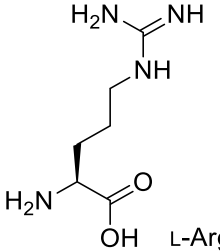 L-Arg | 3 | 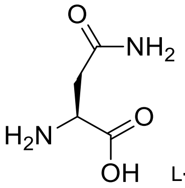 L-Asn |
| 4 | 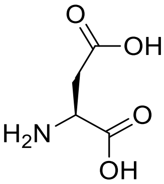 L-Asp | 5 | 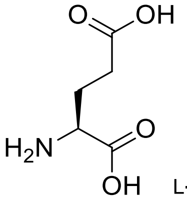 L-Glu | 6 | 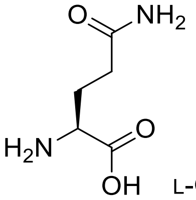 L-Gln |
| 7 | 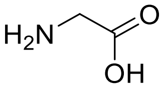 Gly | 8 | 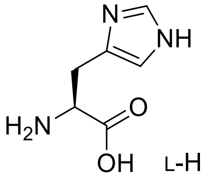 L-His | 9 | 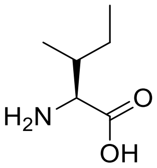 L-Ile |
| 10 | 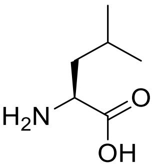 L-Leu | 11 | 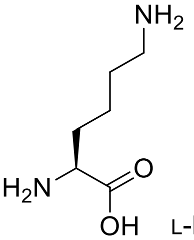 L-Lys | 12 | 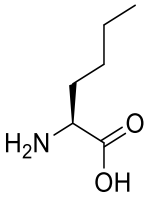 L-Nle |
| 13 | 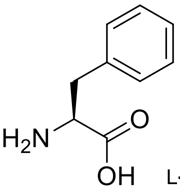 L-Phe | 14 | 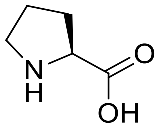 L-Pro | 15 | 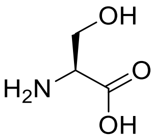 L-Ser |
| 16 | 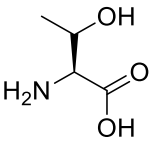 L-Thr | 17 | 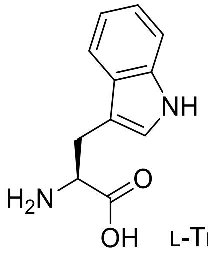 L-Trp | 18 | 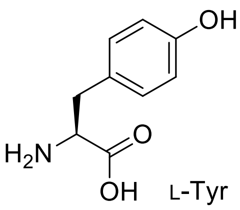 L-Tyr |

19 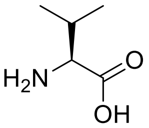 L-Val

20 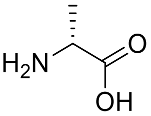 D-Ala

21 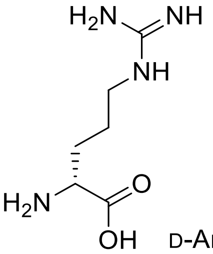 D-Arg

22 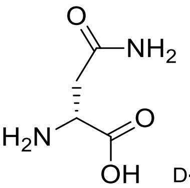 D-Asn

23 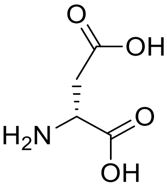 D-Asp

24 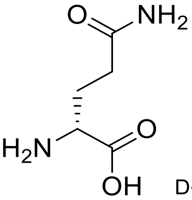 D-Gln

25 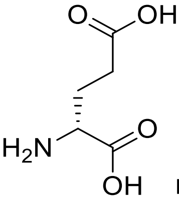 D-Glu

26 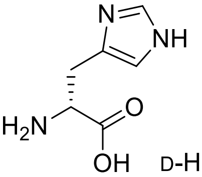 D-His

27 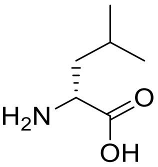 D-Leu

28 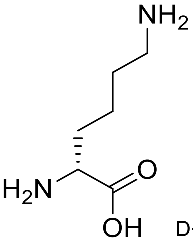 D-Lys

29 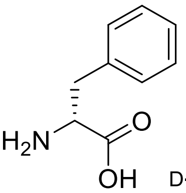 D-Phe

30 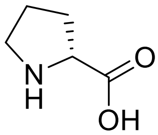 D-Pro

31 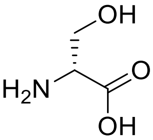 D-Ser

32 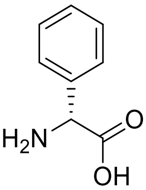 D-Phg

33 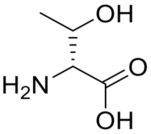 D-Thr

34 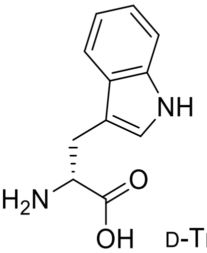 D-Trp

35 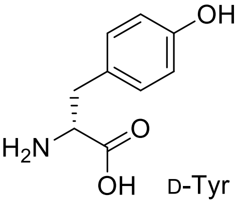 D-Tyr

36 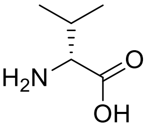 D-Val

37 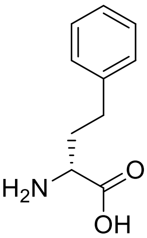 D-hPhe

38 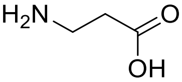 β–Ala

39 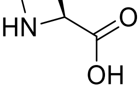 L-Aze

40 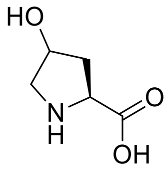 L-Hyp

41 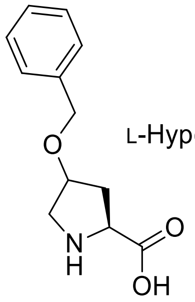 L-Hyp(Bzl)

42 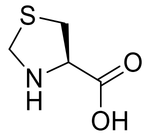 L-Thz

43 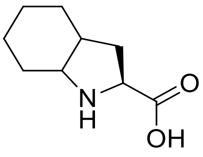 L-Oic

44 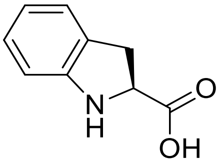 L-Idc

45 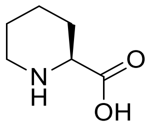 L-Pip

46 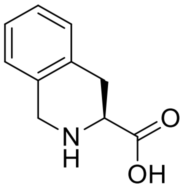 L-Tic

47 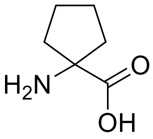 AC5C

48 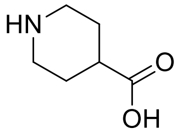 Inp

49 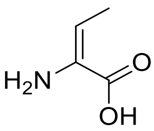 dhAbu

50 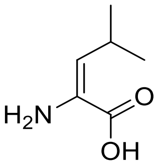 dhLeu

51 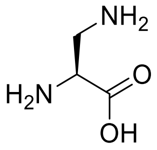 L-Dap

52 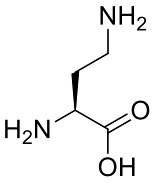 L-Dab

53 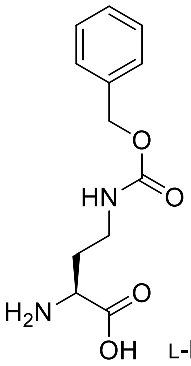 L-Dab(Z)

54 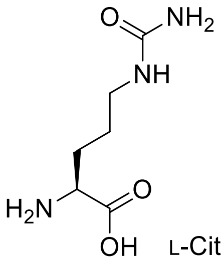 L-Cit

55 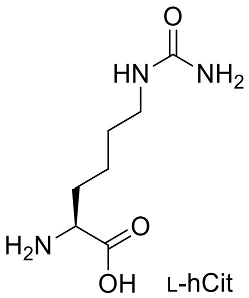 L-hCit

56 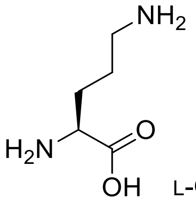 L-Orn

57 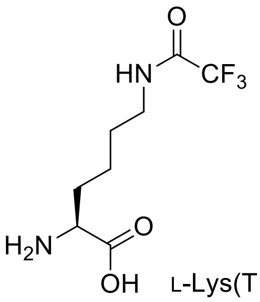 L-Lys(TFA)

58 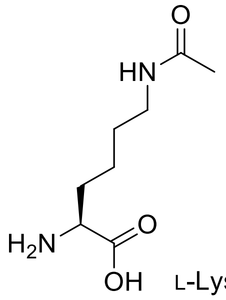 L-Lys(Ac)

59 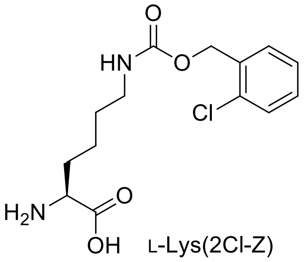 L-Lys(2Cl-Z)

60 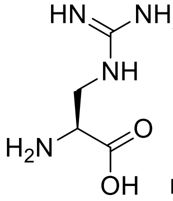 L-Agp

61 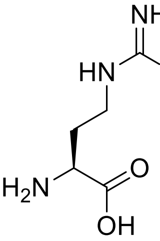 L-Agb

62 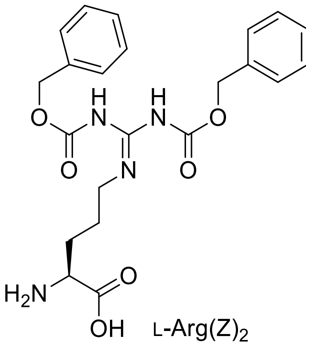 L-Arg(Z)$_2$

63 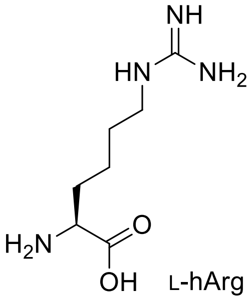 L-hArg

64 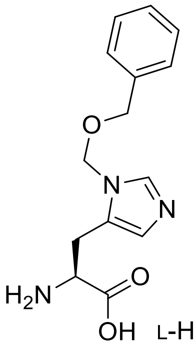 L-His(3-Bom)

65 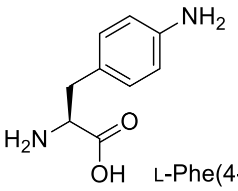 L-Phe(4-NH$_2$)

66 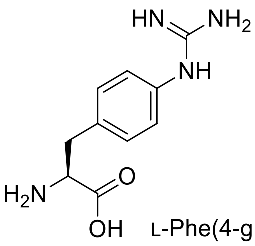 L-Phe(4-guan)

67 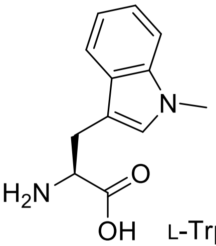 L-Trp(Me)

68 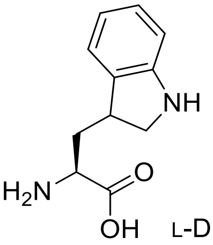 L-Dht

69 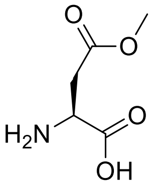 L-Asp(O-Me)

70 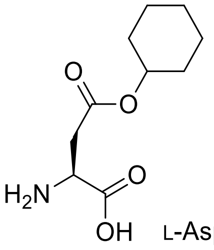 L-Asp(O-cHx)

71 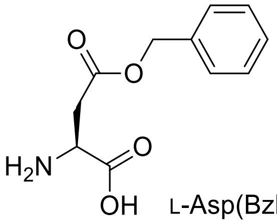 L-Asp(Bzl)

72 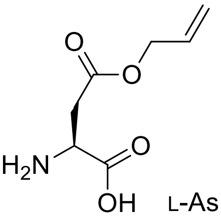 L-Asp(All)

73 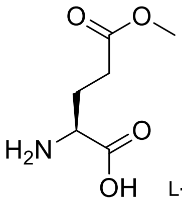 L-Glu(O-Me)

74 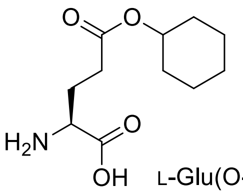 L-Glu(O-cHx)

75 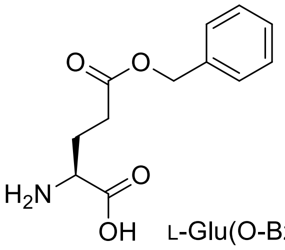 L-Glu(O-Bzl)

76 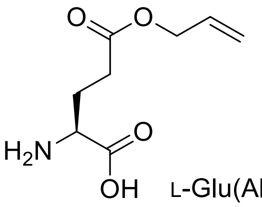 L-Glu(All)

77 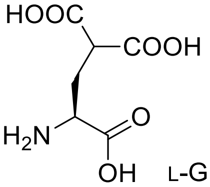 L-Gla

78 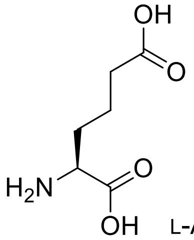 L-Aad

79 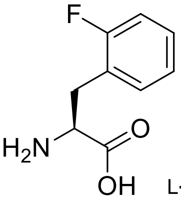 L-Phe(2-F)

80 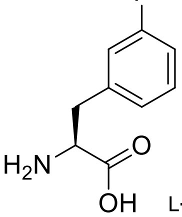 L-Phe(3-F)

81 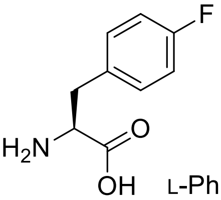 L-Phe(4-F)

82 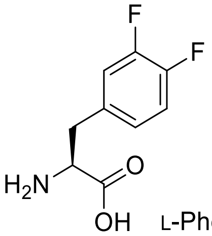 L-Phe(3,4-$F_2$)

83 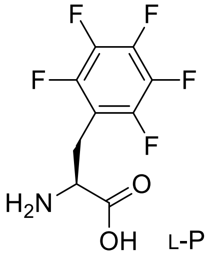 L-Phe($F_5$)

84 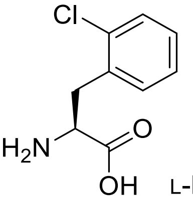 L-Phe(2-Cl)

85 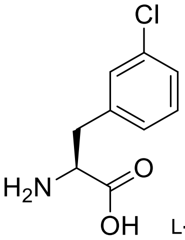 L-Phe(3-Cl)

86 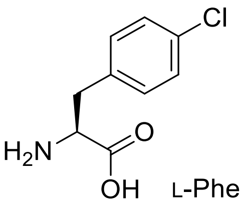 L-Phe(4-Cl)

87 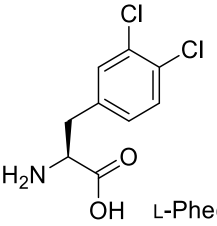 L-Phe(3,4-$Cl_2$)

88 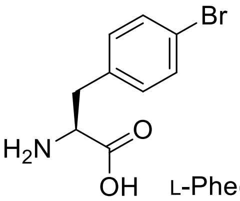 L-Phe(4-Br)

89 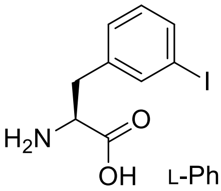 L-Phe(3-I)

90 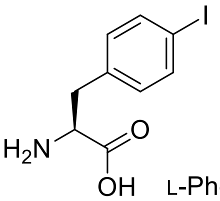 L-Phe(4-I)

91 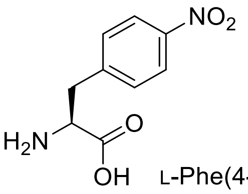 L-Phe(4-$NO_2$)

92 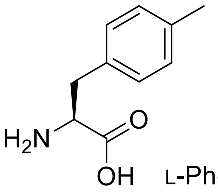 L-Phe(4-Me)

93 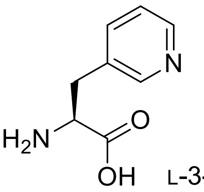 L-3-Pal

94 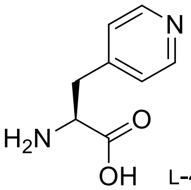 L-4-Pal

95 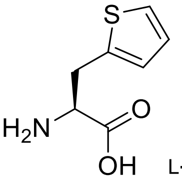 L-Ala(2-th)

96 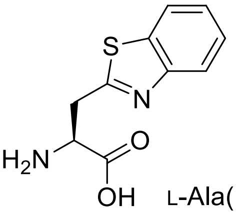 L-Ala(Bth)

97 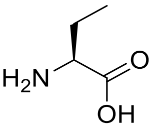 L-Abu

98 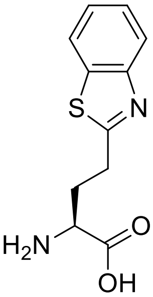 L-Abu(Bth)

99 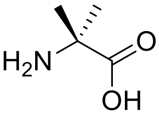 Aib

100 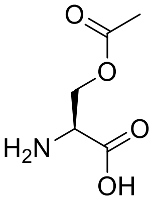 L-Ser(Ac)

101 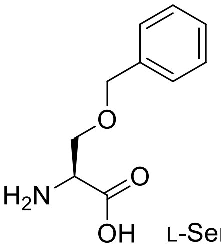 L-Ser(Bzl)

102 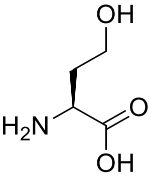 L-hSer

103 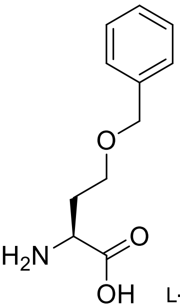 L-hSer(Bzl)

104 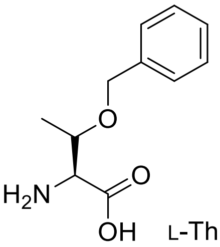 L-Thr(Bzl)

105 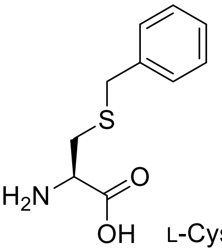 L-Cys(Bzl)

106 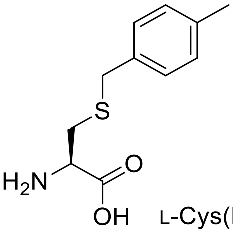 L-Cys(MeBzl)

107 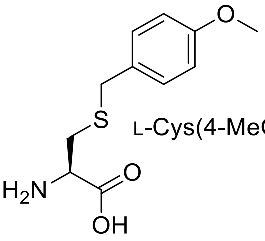 L-Cys(4-MeOBz

108 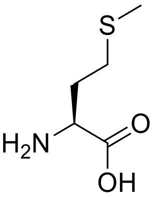 L-Met

109 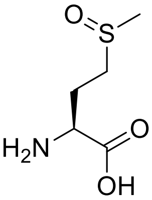 L-Met(O)

110 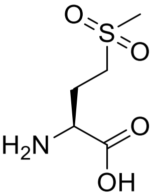 L-Met($O_2$)

111 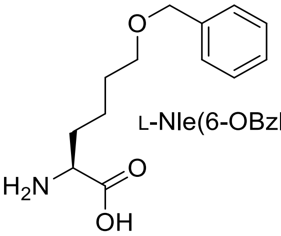 L-Nle(6-OBzl)

112
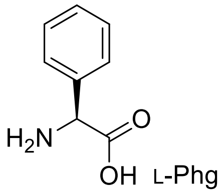


113
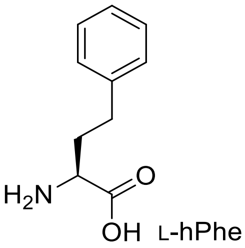


114
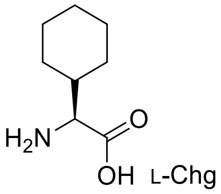


115
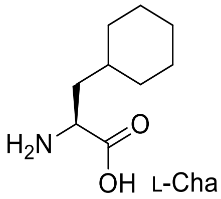


116
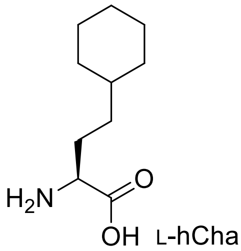


117
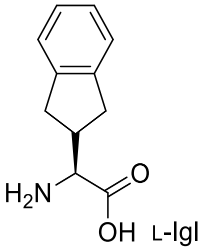


118
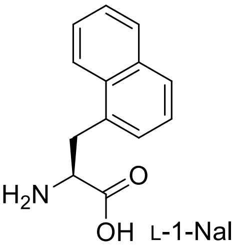


119
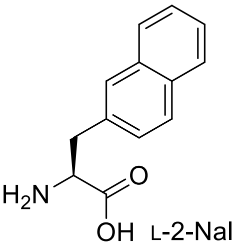


120
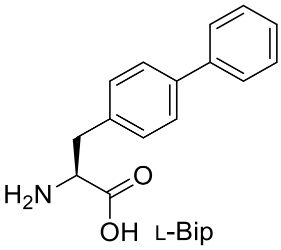


121
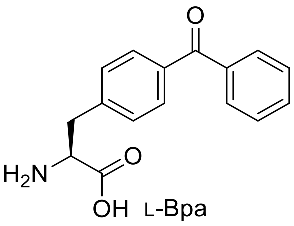


122
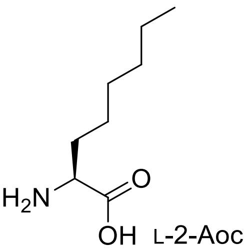


123
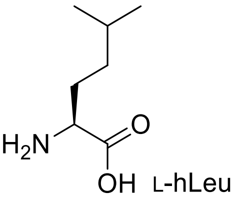


124
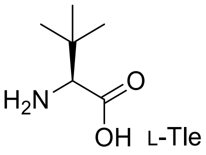


125
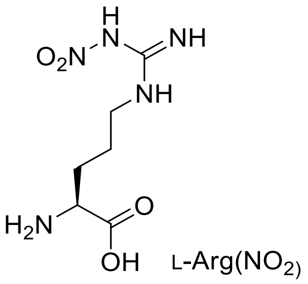


126
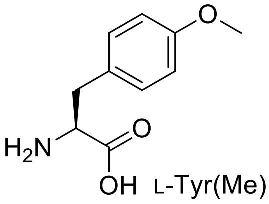


127
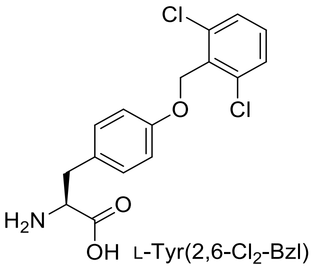


128
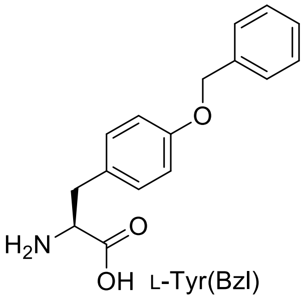


129
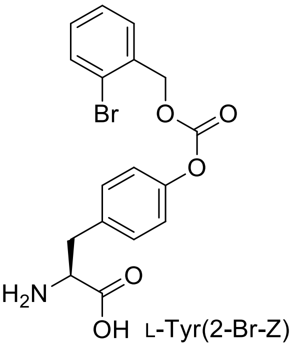

130 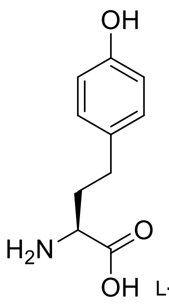 L-hTyr

131 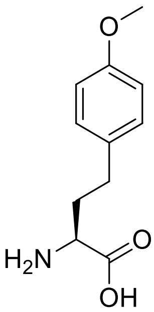 L-hTyr(Me)

132 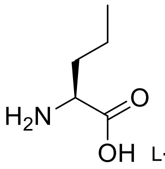 L-Nva

133 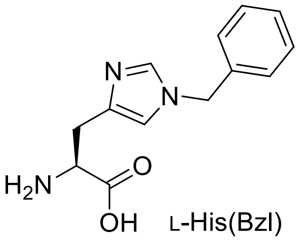 L-His(Bzl)

134 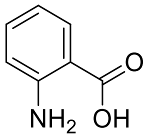 2-Abz

135 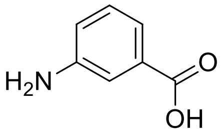 3-Abz

136 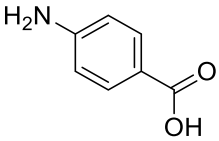 4-Abz

137 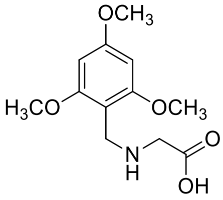 TmbGly

**Table S2.** Crystal diffraction data and model refinement statistics.

| **Protein / Ligand** | **SARS-CoV-2 M$^{pro}$ in complex with biotin-PEG(4)-Abu-Tle-Leu-Gln-vinylsulfone** |
|---|---|
| **PDB entry** | 6Z2E |
| ***Data collection statistics*** | |
| X-ray source | DESY PETRA III P11 |
| Wavelength [Å] | 1.0332 |
| $V_m$ [Å$^3$/Da] | 2.11 |
| Solvent content [%] | 41.77 |
| Space group | $P6_122$ |
| Unit cell dimensions [Å] | $a$ = 104.08, $b$ = 104.08, $c$ = 91.25 |
| Unit cell dimensions [º] | α = β = 90.0, γ = 120.0 |
| Resolution range [a] [Å] | 45.62 - 1.70 (1.79 - 1.70) |
| Number of observations | 1,255,508 (177,717) |
| Number of unique reflections | 32,640 (4,679) |
| Completeness [%] | 100.0 (100.0) |
| Mean I/σ(I) | 32.4 (2.8) |
| Multiplicity | 38.5 (38.0) |
| $R_{merge}$ [b] [%] | 0.074 (1.574) |
| $R_{pim}$ [c] [%] | 0.012 (0.257) |
| $CC_{1/2}$ [d] | 1.000 (0.861) |
| Wilson $B$-factor [Å$^2$] | 29 |
| ***Refinement statistics*** | |

| | |
|---|---|
| Number of unique reflections used for refinement | 32,640 (4,679) |
| $R_{cryst}$ [e] /$R_{free}$ [f] [%] | 19.20/24.34 |
| r.m.s.d. in bond lengths [Å] | 0.01 |
| r.m.s.d. in bond angles [°] | 1.8 |
| Clashscore [g] | 3 |
| Average B-factor for protein atoms [$Å^2$] | 34 |
| Average B-factor for ligand atoms [$Å^2$] | 57 |
| Average B-factor for water molecules [$Å^2$] | 43 |
| Number of protein atoms | 2369 |
| Number of ligand atoms | 70 |
| Number of water molecules | 232 |
| **Ramachandran plot** | |
| Preferred regions [%] | 97.0 |
| Allowed regions [%] | 3.0 |
| Outlier regions [%] | 0.0 |

[a] The highest resolution shell is shown in parantheses.

[b] $R_{merge} = \sum_{hkl} \sum_{i=1}^{n} |I_i(hkl) - \bar{I}(hkl)| / \sum_{hkl} \sum_{i=1}^{n} I_i(hkl)$

[c]$R_{pim} = \sum_{hkl} \sqrt{1/(n-1)} \sum_{i=1}^{n} |I_i(hkl) - \bar{I}(hkl)| / \sum_{hkl} \sum_{i=1}^{n} I_i(hkl)$ 1

[d] $CC_{1/2}$ is the correlation coefficient determined by two random half data sets[2]

[e] $R_{cryst} = \sum_{hkl} |F_o(hkl) - F_c(hkl)| / \sum_{hkl} |F_o(hkl)|$

[f] $R_{free}$ was calculated for a test set of reflections (5%) omitted from the refinement.

[g] Clashscore is defined as the number of clashes calculated for the model per 1000 atoms (including hydrogens) of the model. Hydrogens were added by MolProbity.[3]

### HRMS and analytical chromatograms of synthesized substrates

### Ac-Abu-Tle-Leu-Gln-ACC

m/z: $[M+H]^+$ for $C_{34}H_{49}N_7O_9$ ($m/z_{calcd}$ = 700.3665; $m/z_{found}$ = 700.3669)

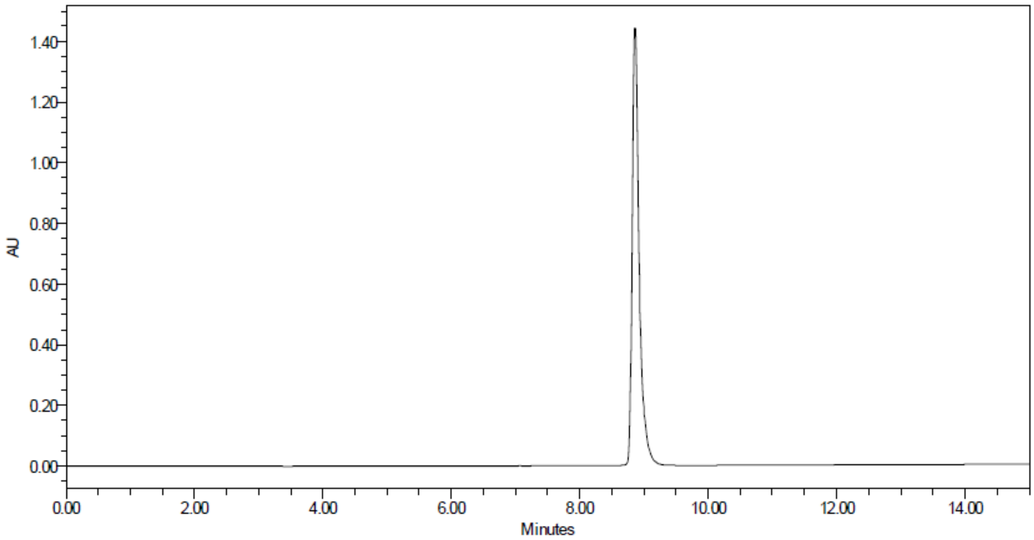


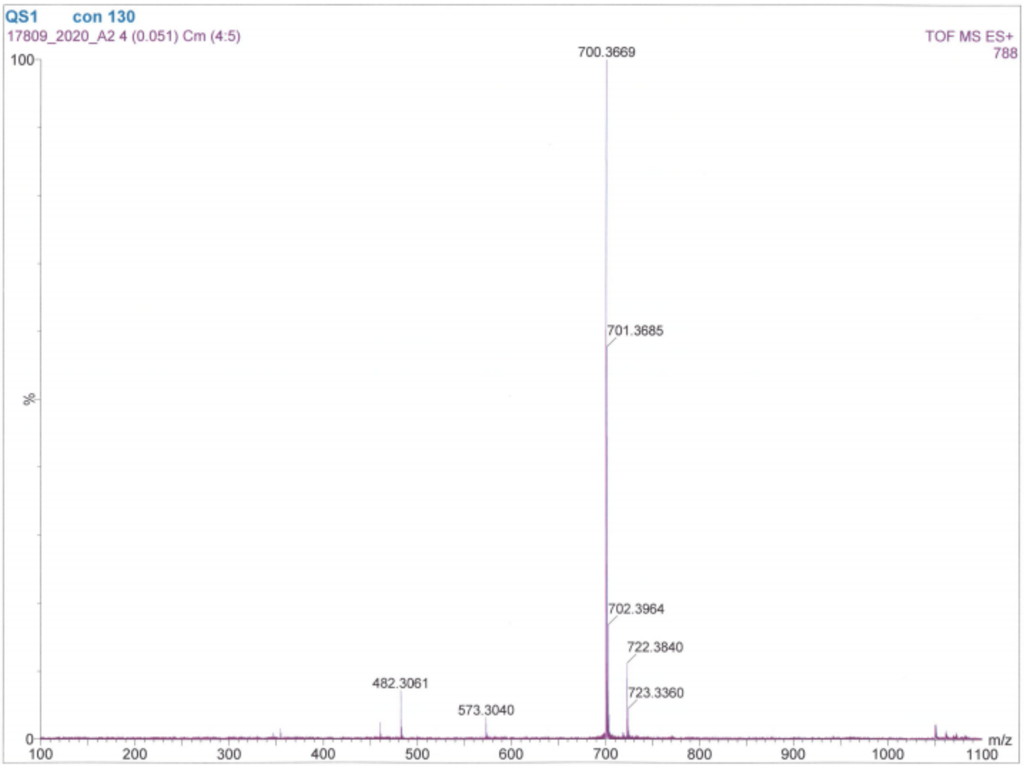

**Ac-Val-Tle-Leu-Gln-ACC**

m/z: $[M+H]^+$ for $C_{35}H_{52}N_8O_9$ ($m/z_{calcd}$ = 729.3931; $m/z_{found}$ = 729.3934)

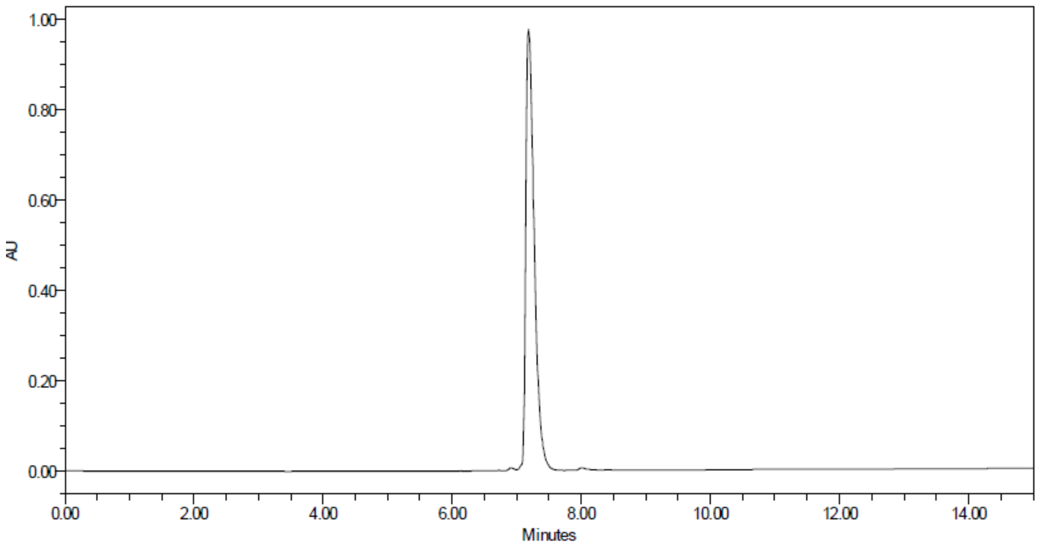


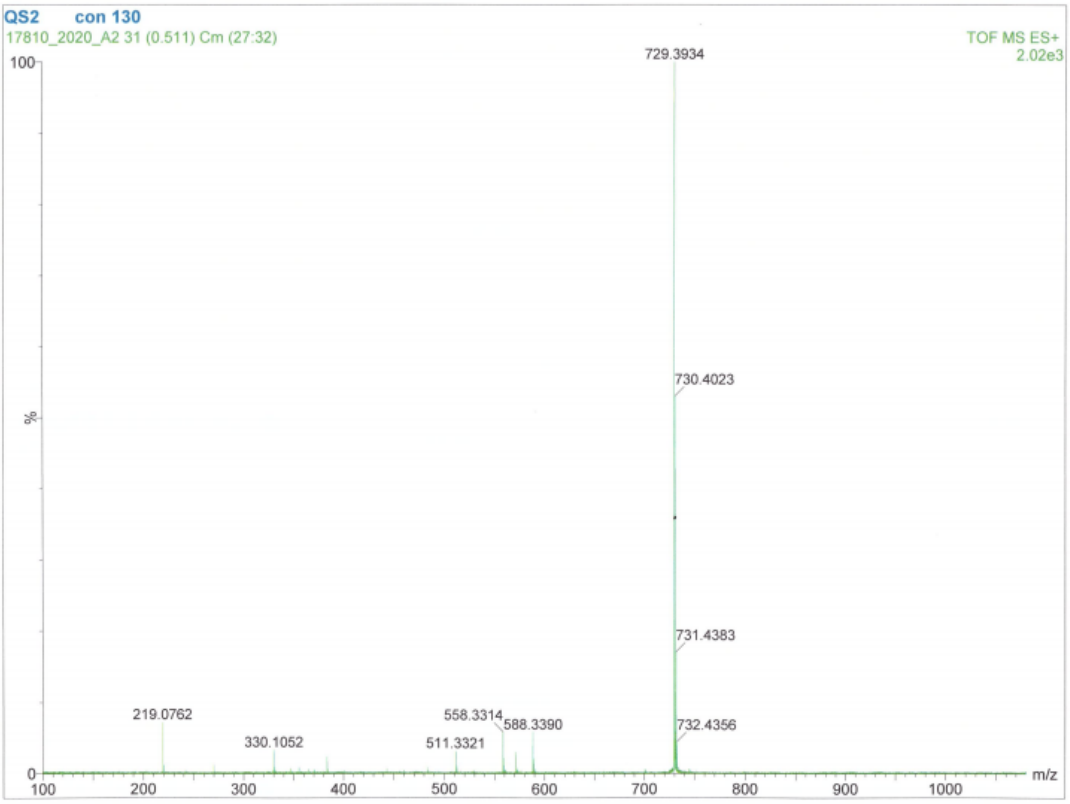

**Ac-Ala-Tle-Leu-Gln-ACC**

m/z: $[M+H]^+$ for $C_{33}H_{48}N_8O_9$ ($m/z_{calcd}$ = 701.3618; $m/z_{found}$ = 701.3620)

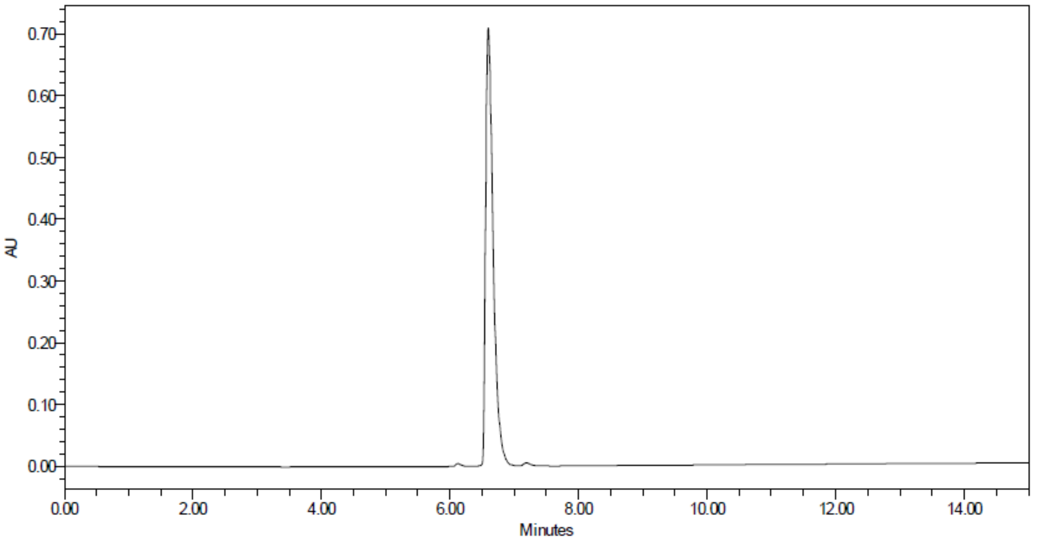


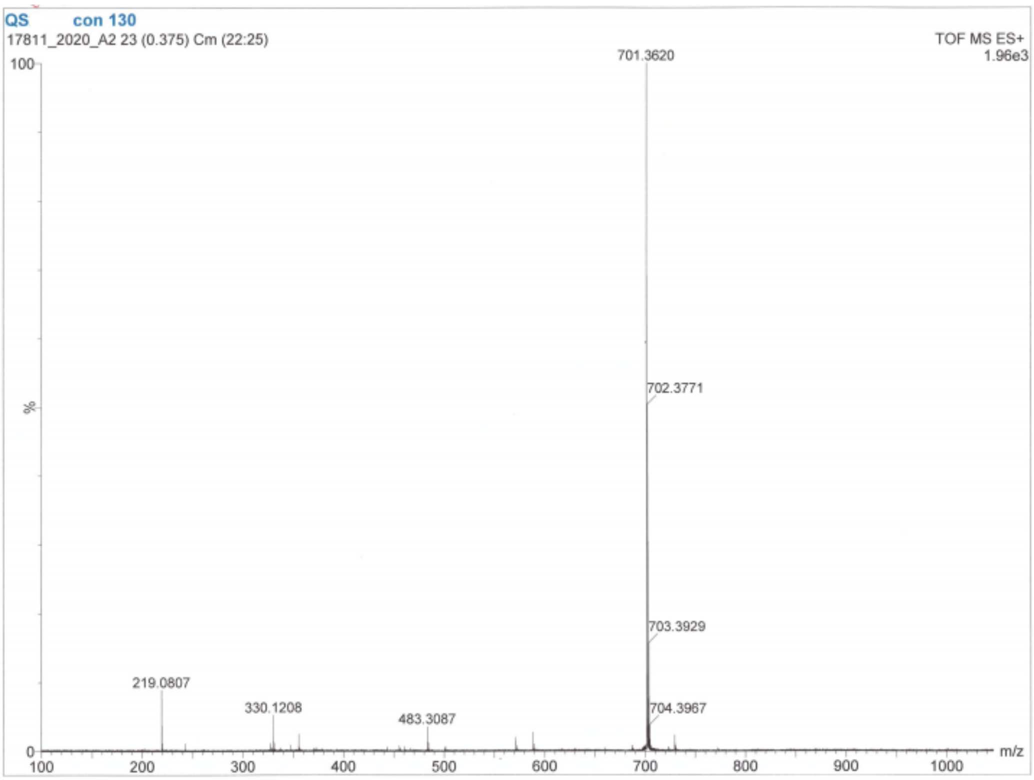

**Ac-Thz-Tle-Leu-Gln-ACC**

m/z: $[M+H]^+$ for $C_{34}H_{47}N_7O_9S$ ($m/z_{calcd}$ = 730.3229; $m/z_{found}$ = 730.3237)

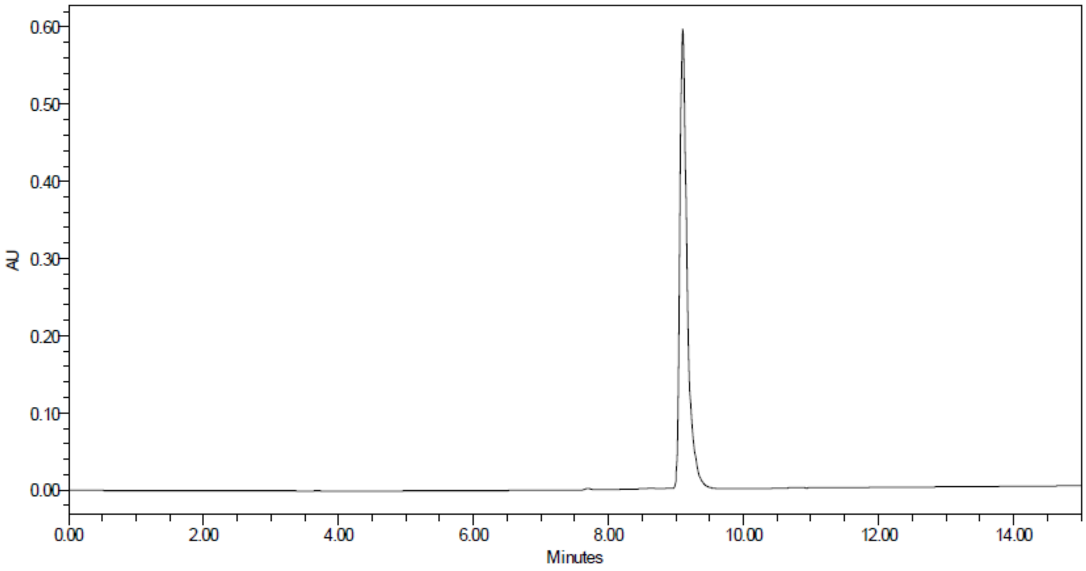


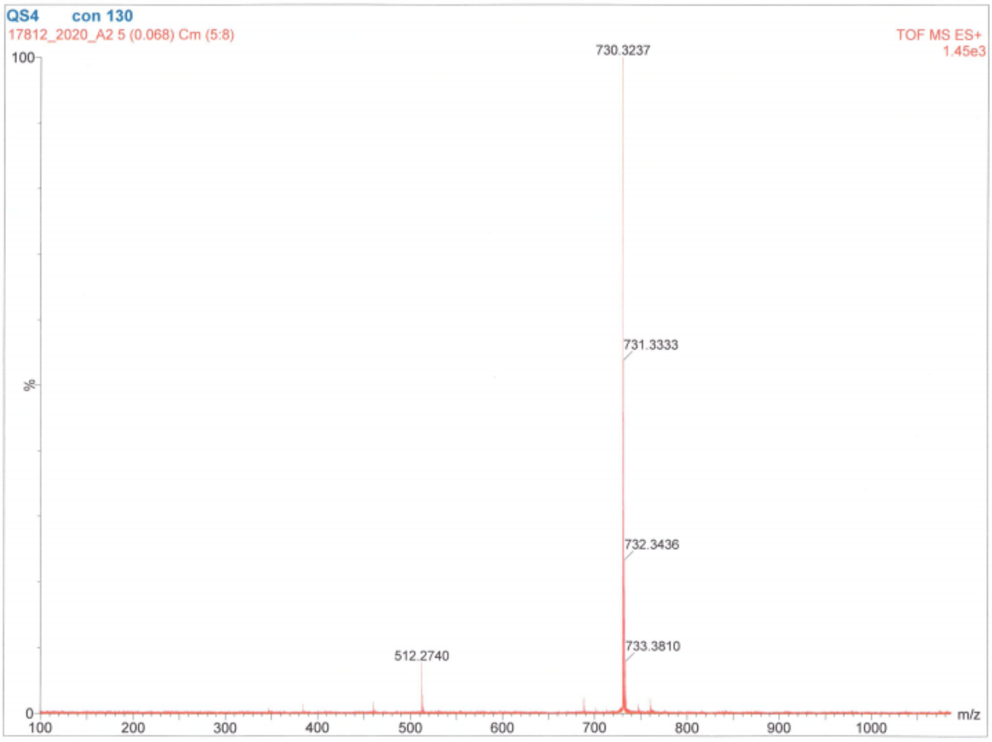

**Ac-Abu-DTyr-Leu-Gln-ACC**

m/z: [M+H]+ for $C_{37}H_{47}N_7O_{10}$ ($m/z_{calcd}$ = 750.3458; $m/z_{found}$ = 750.3453)

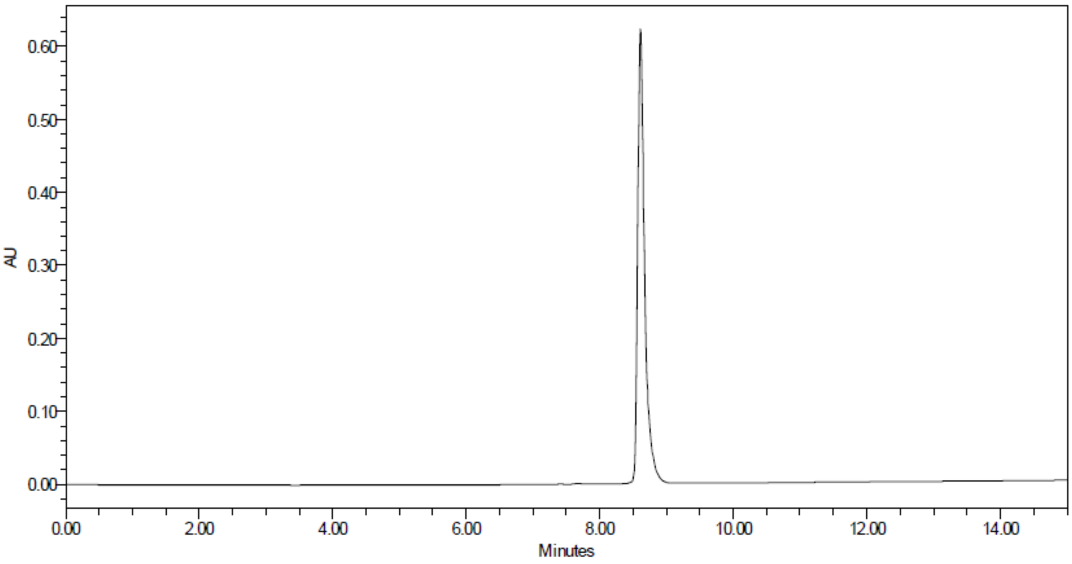


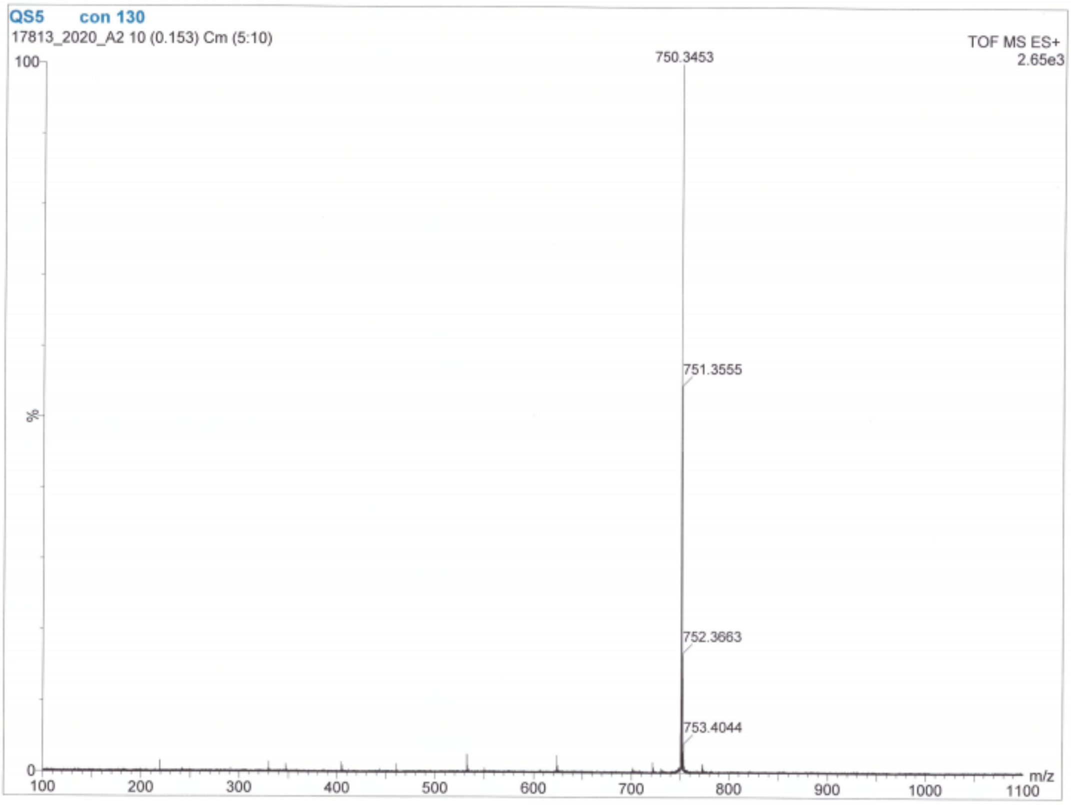

**Ac-Abu-Orn-Leu-Gln-ACC**

m/z: $[M+H]^+$ for $C_{33}H_{48}N_8O_9$ ($m/z_{calcd}$ = 701.3618; $m/z_{found}$ = 701.3618)

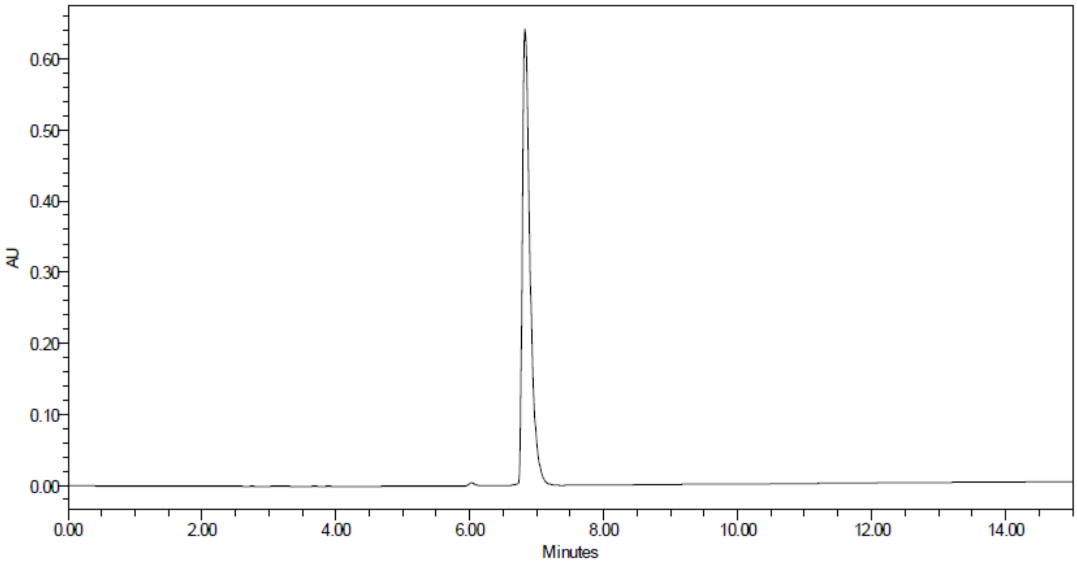


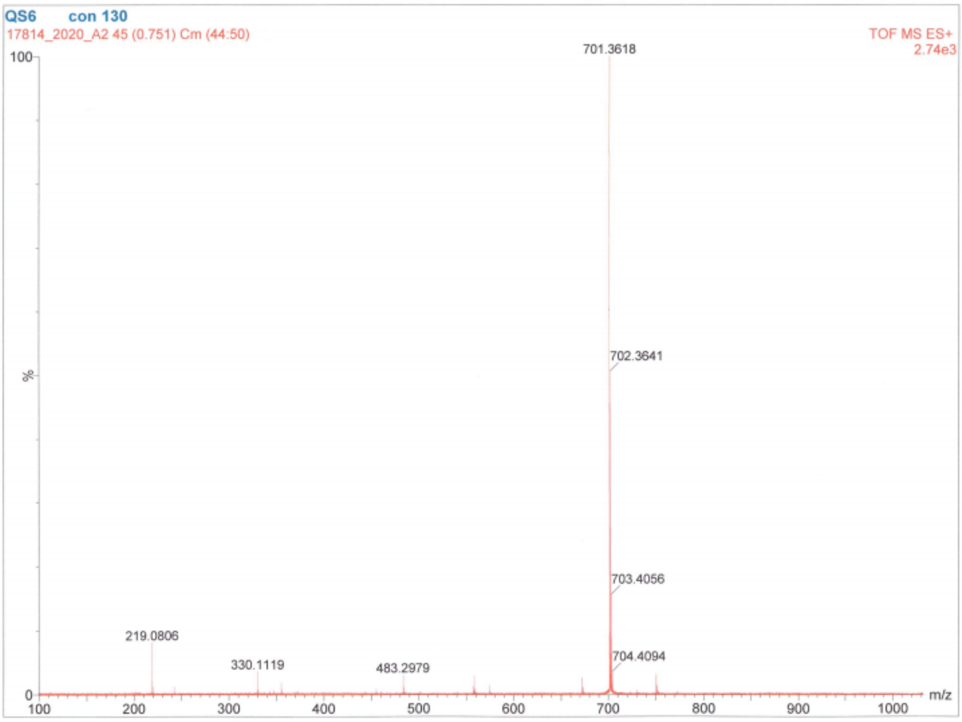

**Ac-Abu-Lys-Leu-Gln-ACC**

m/z: $[M+H]^+$ for $C_{34}H_{50}N_8O_9$ ($m/z_{calcd}$ = 715.3774; $m/z_{found}$ = 715.3779)

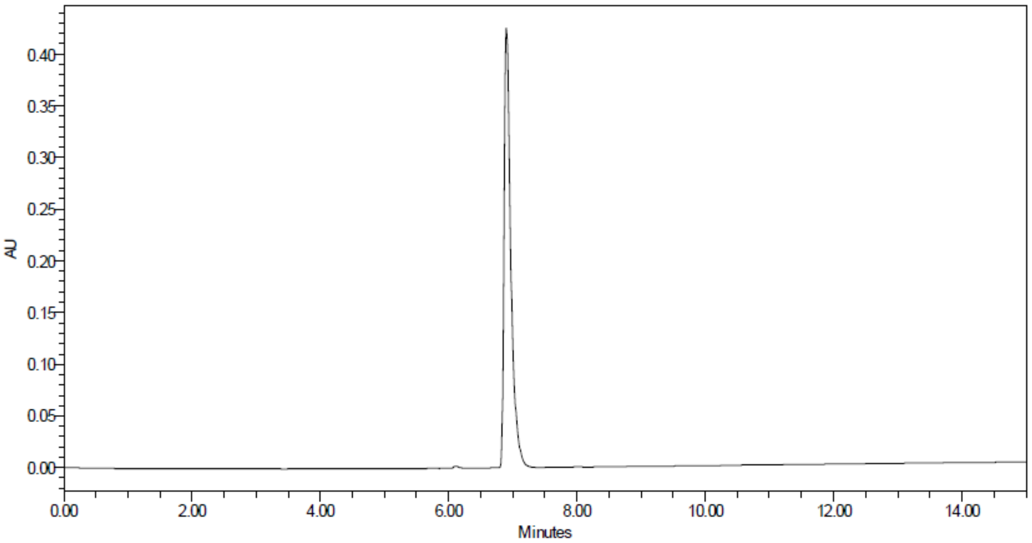


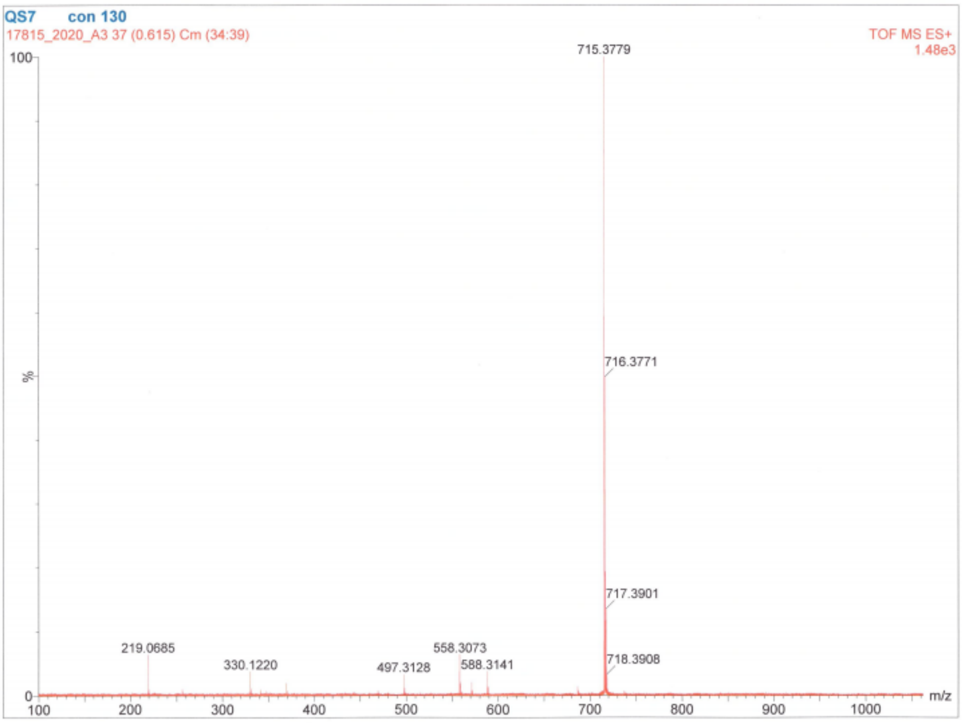

**Ac-Abu-Tle-hLeu-Gln-ACC**

m/z: $[M+H]^+$ for $C_{35}H_{51}N_7O_9$ ($m/z_{calcd}$ = 714.3822; $m/z_{found}$ = 714.3821)

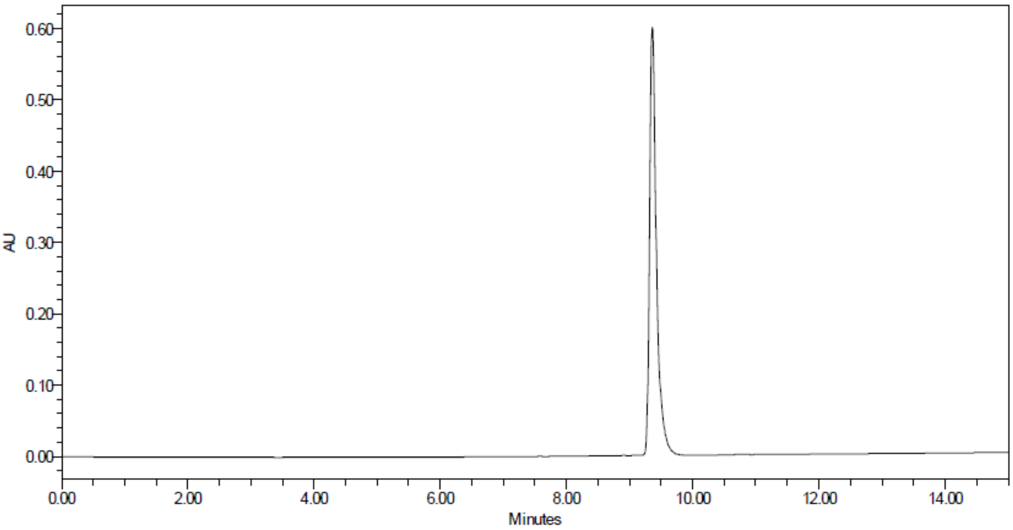


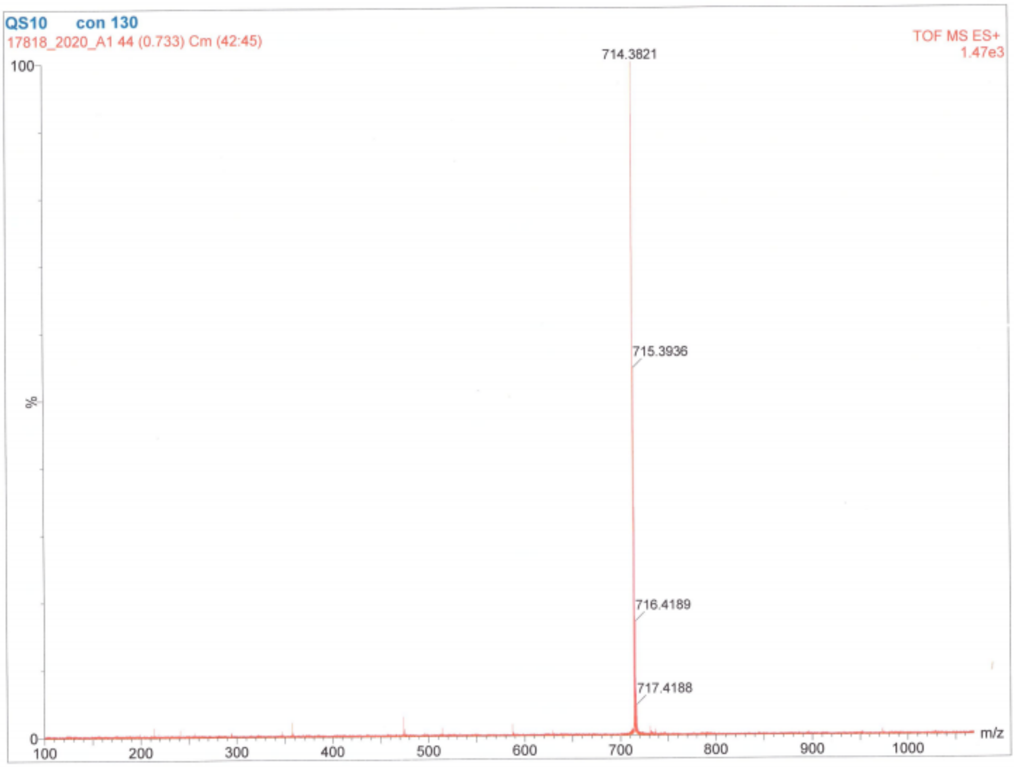

**Ac-Abu-Tle-Leu-Gln-VS**

m/z: $[M+H]^+$ for $C_{25}H_{45}N_5O_7S$ ($m/z_{calcd}$ = 560.31; $m/z_{found}$ = 560.16)

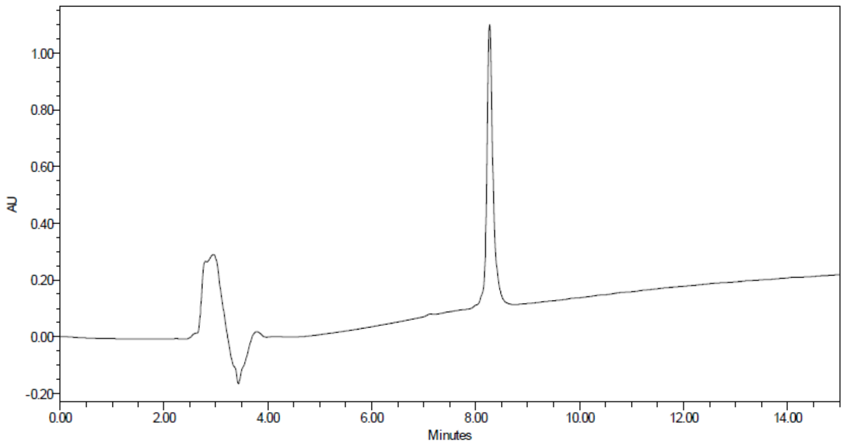


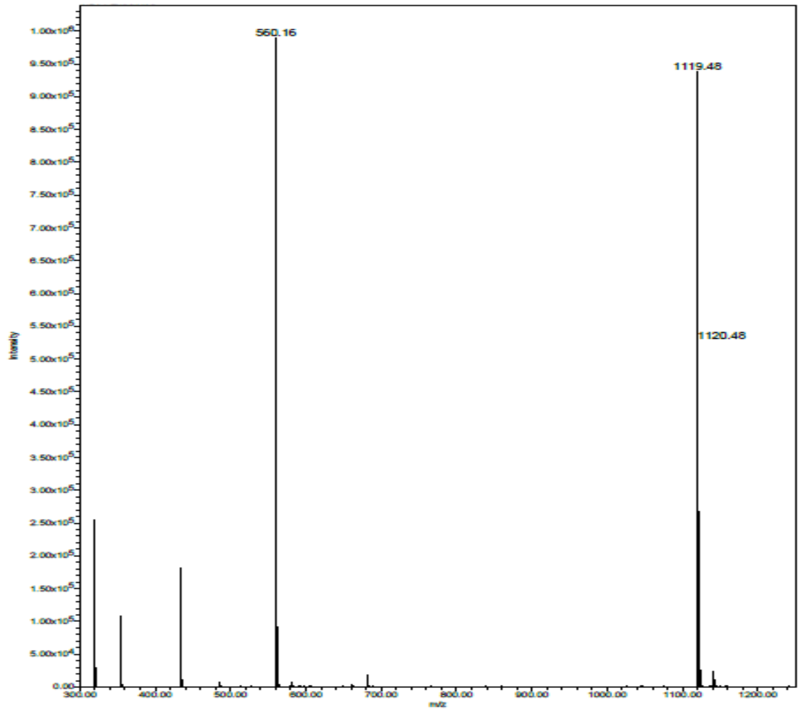

**Biotin-PEG(4)-Abu-Tle-Leu-Gln-VS**

m/z: $[M+H]^+$ for $C_{44}H_{78}N_8O_{13}S_2$ ($m/z_{calcd}$ = 991.52; $m/z_{found}$ = 991.38)

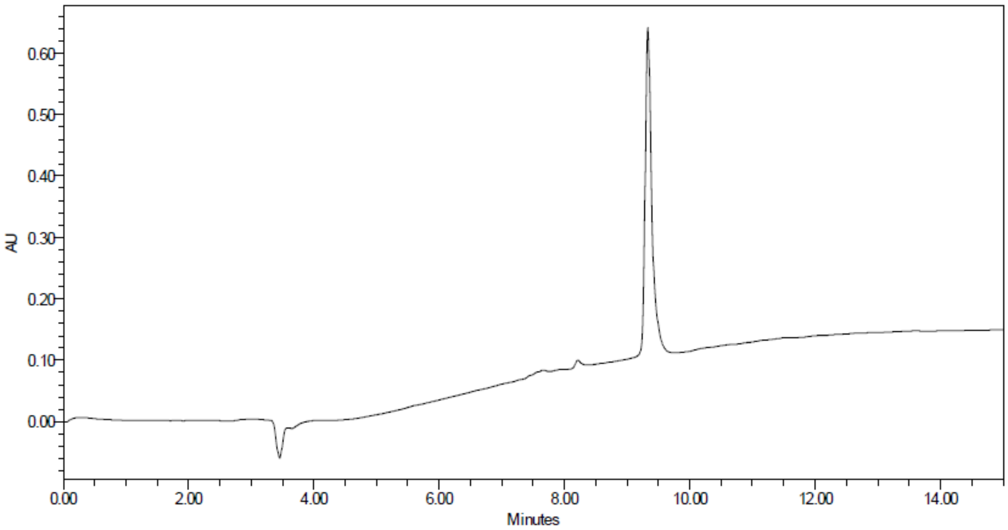


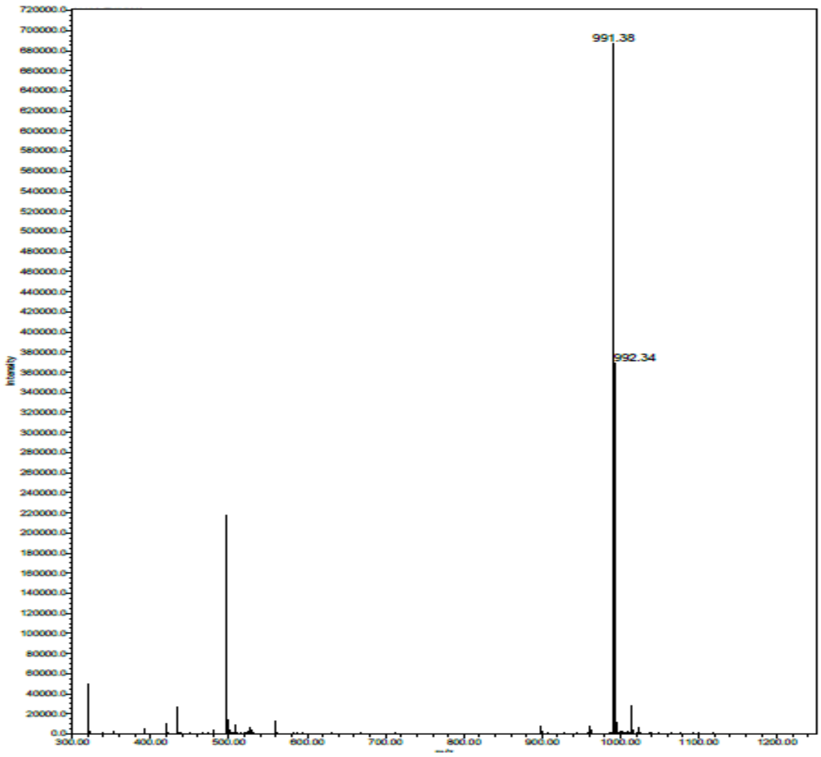

**Cy5-PEG(4)-Abu-Tle-Leu-Gln-VS**

m/z: $[M+H]^+$ for $C_{66}H_{101}N_8O_{12}S$ ($m/z_{calcd}$ = 1229.73; $m/z_{found}$ = 1229.70)

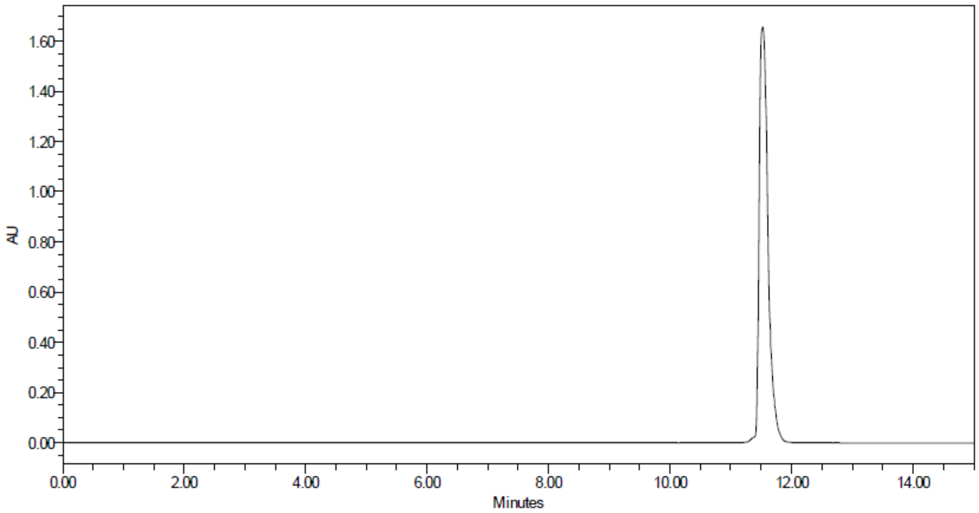


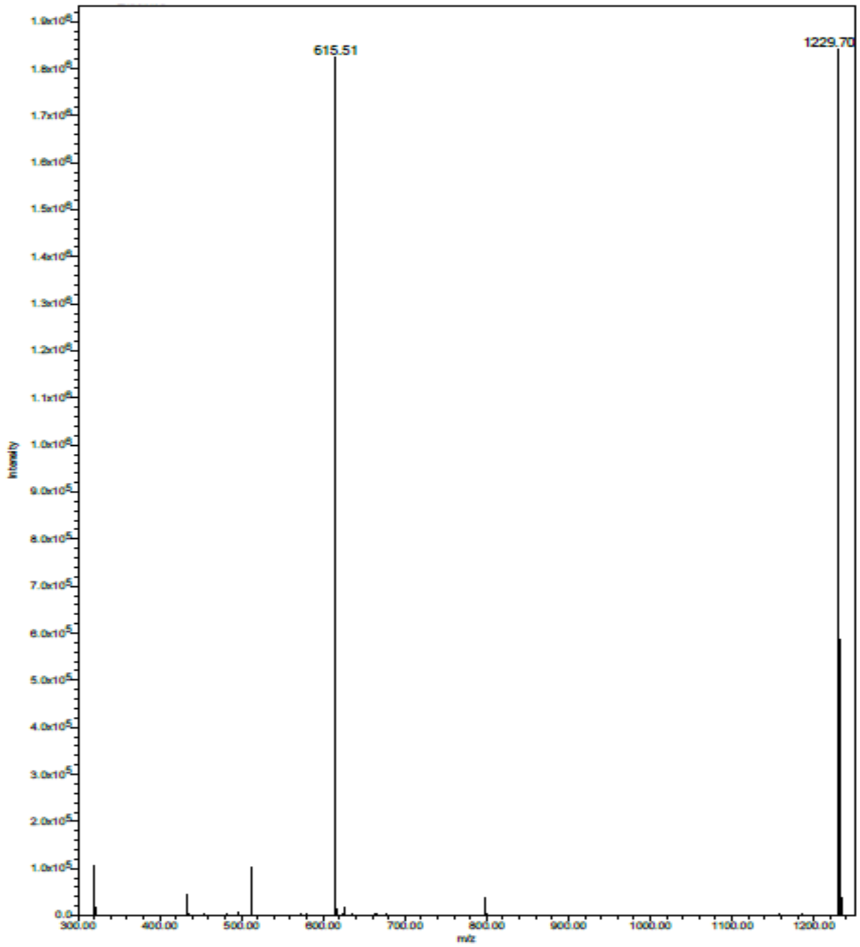

1. Weiss, M. S.; Hilgenfeld, R., On the use of the merging R factor as a quality indicator for Xray data. *J. Appl. Cryst.* **1997**, *30*, 203-5.
2. Karplus, P. A.; Diederichs, K., Linking crystallographic model and data quality. *Science* **2012,** *336* (6084), 1030-3.
3. Chen, V. B.; Arendall, W. B., 3rd; Headd, J. J.; Keedy, D. A.; Immormino, R. M.; Kapral, G. J.; Murray, L. W.; Richardson, J. S.; Richardson, D. C., MolProbity: all-atom structure validation for macromolecular crystallography. *Acta Crystallogr D Biol Crystallogr* **2010,** *66* (Pt 1), 12-21.
